# Confidence Scoring for AI-Predicted Antibody–Antigen Complexes: AntiConf as a Precision-driven Metric

**DOI:** 10.1101/2025.07.25.666870

**Authors:** Serbulent Unsal, Benjamin Holland, Inci Sardag, Emel Timucin

## Abstract

Accurate determination of antibody-antigen (Ab-Ag) complex structures is critical for therapeutic development. While AI-based methods, beginning with AlphaFold2 (AF2), have revolutionized multimer predictions, the optimal strategies for Ab-Ag modeling and the reliability of their confidence scores remain active areas of research. This study evaluates the performance of AF2, Boltz-1, Boltz-1x, Boltz-2, Chai-1, Protenix and ESMFold, on a curated dataset of 200 antibody-antigen (Ab-Ag) complexes. Our findings reveal that AF2 remains a strong predictor against newer AlphaFold3 (AF3)-inspired methods in Ab-Ag complex prediction. Chai-1 consistently ranks as the second-best performer across multiple success metrics. We observed diverse effects of recycling iterations, with AF2, Chai-1 and Protenix benefiting from increased cycles, unlike Boltz variants. We analyzed various model confidence scores, noting high precision from pDockQ2 and high recall from pTM. By integrating these two scores, we developed AntiConf, a novel metric that achieves superior performance for all methods in terms of precision and recall. These strengths make AntiConf a valuable post score for both computational predictions and downstream experimental workflows, reflecting its potential to improve Ab-Ag complex predictions by AF2 and AF3 architectures. Altogether, this study addresses current limitations in AI-based Ab-Ag complex prediction, showcasing the potential of AntiConf for future predictions and assessments of these complexes, and providing a guideline for improving the accuracy of Ab-Ag complex prediction.

## Introduction

Antibodies (Abs) are central to adaptive immunity, functioning either as neutralizing agents and flagging antigens for clearance. Their therapeutic importance has grown considerably due to their involvement in a wide spectrum of successful immunotherapy applications in cancer, autoimmune disorders, and infectious diseases, transforming the landscape of modern medicine [1].

Structurally, antibodies are Y-shaped proteins composed of two identical heavy and two identical light chains stabilized by disulfide bonds. Monoclonal antibodies (mAbs) consist of paired heavy and light chains, whereas nanobodies (Nbs) are composed of a single variable heavy chain domain. In both mAbs and Nbs, antigen recognition is primarily mediated by three hypervariable loops known as the complementarity-determining regions (CDRs), collectively forming the paratope region [2–5]. Given the smaller size and extended CDR loops, Nbs can access epitopes often unreachable by mAbs [6, 7]. Among CDRs, the CDR H3 loop is the most diverse region in terms of structure, length and amino acid composition, and forms the highest number of antigen contacts [8–13]. The length and structure of the CDR H3 loop is known to affect the orientation and engagement of neighboring CDRs, thereby modulating the overall paratope architecture and specificity [8].

Experimental determination of multimer structures including Ab-Ag complexes is costly and time-intensive, underscoring the importance of computational methods as cost-effective alternatives to experiments [14]. Despite the growing availability of tools for designing antibodies and nanobodies, accurate prediction of antibody-antigen complex structures remains a major challenge [15–17]. Essentially, accurate modeling of the CDR H3 loop continues to pose significant difficulties [18, 19], making it a persistent bottleneck in generating high-resolution antibody-antigen complex structures [20].

Recent developments in deep learning-based structure prediction methods have raised the bar for computationally predicted multimeric structures. Essentially, AlphaFold2 (AF2) has shown striking success in predicting protein-protein complexes [21, 22]. AlphaFold3 (AF3) introduces a diffusion-based architecture [23] and has demonstrated improved accuracy over AF2 for predicting multimeric structures [23]. In parallel with the development of AF3, several AF3-inspired open-source models have emerged, each addressing distinct limitations of the original framework. Among these, Boltz-1 implements a deep learning model refining the AF3 framework with improved input processing, architectural adjustments, and a redesigned confidence model [24]. The recent version Boltz-2 represents an advancement over Boltz-1 through its ability to predict binding affinity between small molecules and proteins, representing a leap from predicting static structures to understanding dynamic molecular interactions [25]. Chai-1, another AF3-inspired model with the same model architecture and training strategy with AF3, supports experimental restraints for enhanced accuracy and can function without MSAs [26]. Protenix, another refined reproduction of AF3, incorporates clarification and correction of ambiguous steps, typographical errors, and targeted adjustments informed by model behavior [27]. In addition to these AF3-inspired models, other approaches are also available such as ESMFold, which uses a transformer-based architecture bypassing multiple sequence alignment (MSA) generation [28]. While all these methods represent important steps forward in protein-protein complex prediction, their effectiveness for Ab-Ag systems needs further evaluations to identify where they succeed, and where performance gaps exist.

To this end, we assessed the performance of AF2 [21] and AF3-based methods including Boltz-1 [24], Boltz-1x (Boltz-steering) [24], Boltz-2 [25], Chai-1 [26], Protenix [27] along with ESMFold [28], on a curated set of antibody-antigen complexes that are absent from the training data of these methods. For each complex, we generated 25 predictions and extensively evaluated their performance based on the scores including but not limited to ipTM, ipLDDT, iPAE, and pDockQ2. We introduced a novel score, AntiConf, developed by integrating pTM and pDockQ2, which consistently outperformed all evaluated metrics in terms of both precision and recall. Our findings highlight the strengths and limitations of AF2- and AF3-based approaches in modeling real-world antibody–antigen complexes, demonstrating AntiConf’s potential for improving Ab-Ag complex predictions by AF2 and/or AF3-based methods.

## Methods

### Dataset Construction

We assembled a curated dataset of 200 Ab–Ag complexes by combining and refining data from two recent benchmarking studies [29, 30]. The first subset, consisting of 39 complexes from Yin et al. (2024) [29], was directly incorporated. The second subset was derived from the 210 complexes compiled by Hitawala et al. (2024) [30], after filtering out unbound antibody structures and complexes involving non-proteinaceous antigens. Additionally, we corrected metadata discrepancies in the original dataset, particularly those related to missing or unmodeled regions in the input sequences. Redundant entries representing alternative biological units of the same PDB target were also removed, retaining only one representative per complex. The final dataset comprises 200 Ab–Ag complexes, including structures released after the training cutoff dates of AF2 and other prediction methods. All selected sequences were verified to have low sequence identity to training data, ensuring the inclusion of previously unseen antibody–antigen interactions.

### Complex Predictions

AF2 v2.3.2 was implemented through ColabFold [22, 31] using MMseqs2 [32] and the UniRef+Environmental database [33]. To generate a total of 25 predictions per complex, we used a different random seed for each prediction, producing five predictions for each AF2 prediction. Boltz-1 and Boltz-1x [24], Boltz-2 [25], Chai-1 [26] and Protenix [27] were used to generate 25 models for each target using a single seed. Before assembling the final model, different number of recycling iterations were tested for all methods except for ESMFold. For ESMFold, only a single model was generated for each target [28]. All predictions were conducted via Tamarind Bio [34]. The dataset and predicted models for all methods are available at https://doi.org/10.5281/zenodo.16404871.

### Evaluation of Complex Model Accuracy

Model accuracy was assessed by DockQ v2 [35, 36] and Critical Assessment of Prediction of Interactions (CAPRI) classification system [37, 38]. CAPRI classification system groups models into discrete categories such as incorrect, acceptable, medium, and high using fixed cutoff values of three complex quality indicators of fraction of native contacts (*f*_*nat*_), ligand backbone RMSD (lRMSD), and interface backbone RMSD (iRMSD). DockQ provides a continuous score that is calculated by combining these three indicators. Fixed DockQ thresholds, i.e 0.23, 0.49, 0.80, are also used to align with the respective CAPRI categories.

### Calculation of Interface pLDDT and PAE

For each chain in the structure, a residue was classified as an interface residue if the distance between its C*α* atom and the C*α* atom of any residue from a neighboring chain was less than or equal to the defined threshold of 8 Å Only C*α*–C*α* distances were considered to ensure consistency across all residue types. For each identified interface residue, the corresponding pLDDT score was extracted from the model files. PAE scores for each interacting chain pair were obtained from the corresponding interface region in the global PAE matrix. A normalized interface PAE score was computed using the formula from the ref. [39]. The ipLDDT and iPAE scores were computed for each chain and reported as the average for either the antigen alone or for all chains in the complex.

### Calculation of pDockQ2 and pTMDockQ2

We calculated the pDockQ2 score, which combines ipLDDT and normalized iPAE. Specifically, the product of ipLDDT and normalized iPAE scores was fitted to a four-parameter sigmoid function to obtain the pDockQ2 score [39]:

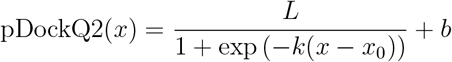

where *x* represents the product of ipLDDT and normalized iPAE. The curve parameters were adopted from ref. [39] as follows: *L* = 1.3103, *x*_0_ = 84.7326, *k* = 0.0747, and *b* = 0.0050. Interface pLDDT, PAE and pDockQ2 calculations were performed using the `pdockq2.py` script available at https://gitlab.com/ElofssonLab/afm-benchmark.

A composite score was computed as a weighted linear combination of pTM and pDockQ2 scores as follows:

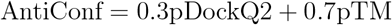

### Statistical Analysis

All statistical analyses were performed using Python libraries and SciPy packages. To evaluate classification performance, we calculated precision, recall, specificity, F1 score, and the area under precision-recall curve. Linear relationships between DockQ score and model metrics were assessed using the Pearson correlation. All reported statistical tests were two-sided, and significance was assessed at a threshold of 0.05.

## Results and Discussion

### Construction and Prediction of the Ab-Ag Complex Dataset

We utilized a curated dataset of 200 antibody-antigen (Ab-Ag) complexes, building upon two prior benchmarking studies [29, 30]. This involved directly incorporating 39 complexes from one source [29] and refining the initial set of 210 Ab-Ag complexes from another source [30] by removing unbound antibody structures, non-proteinaceous antigens, and correcting for missing PDB regions. Redundant sequences, representing different biological units of the same PDB target, were also removed to ensure a unique biological unit per target. The final 200-complex dataset includes structures released after the training cutoffs of methods and thus features novel Ab-Ag interactions with minimal sequence identity to Ab-Ag complexes in the training data of the methods.

We predicted these Ab-Ag complexes using seven different methods; AF2 (v2.3.2) [21], Boltz-1 [24], Boltz-1x (Boltz-steering) [24], Boltz-2 [25], Chai-1 [26], Protenix [27] and ESMFold [28]. Among these methods, Boltz-2 was evaluated on a subset of 18 complexes with deposition dates after its 2023 training cutoff to ensure fair assessment. For Protenix, predictions failed on 26 large complexes, limiting its evaluation to 174 cases. All other methods were successfully run on the full set of 200 Ab-Ag complexes.

We initially examined the concordance between DockQ scores [35, 36] and CAPRI classification [38, 40]. Fig. S1 illustrate the distribution of DockQ scores across CAPRI categories for each method. While predictions showed a clear separation of DockQ among the CAPRI categories with progressively increasing DockQ scores from incorrect to high quality CAPRI classes for each method, some overlaps exist mostly between incorrect-to-acceptable and acceptable-to-medium transitions for all methods. Overall, we found that DockQ scores and thresholds are robust and generally align well with the respective CAPRI categories. We only highlighted DockQ’s limitation for models whose scores lie near its classification boundaries.

### Performance Assessment Based on CAPRI Classification

Fig. 1 summarizes the success rates of six different structure prediction methods, AF2, Boltz1, Boltz-1x, Boltz-2, Chai-1, and Protenix for overall complex set, for antibody-antigen (Ab) interactions, and for nanobody-antigen (Nb) interactions. For each case, we reported success rates based on the number of correctly predicted unique targets achieving at least acceptable accuracy, as well as those falling within each CAPRI category.

**Fig. 1.**
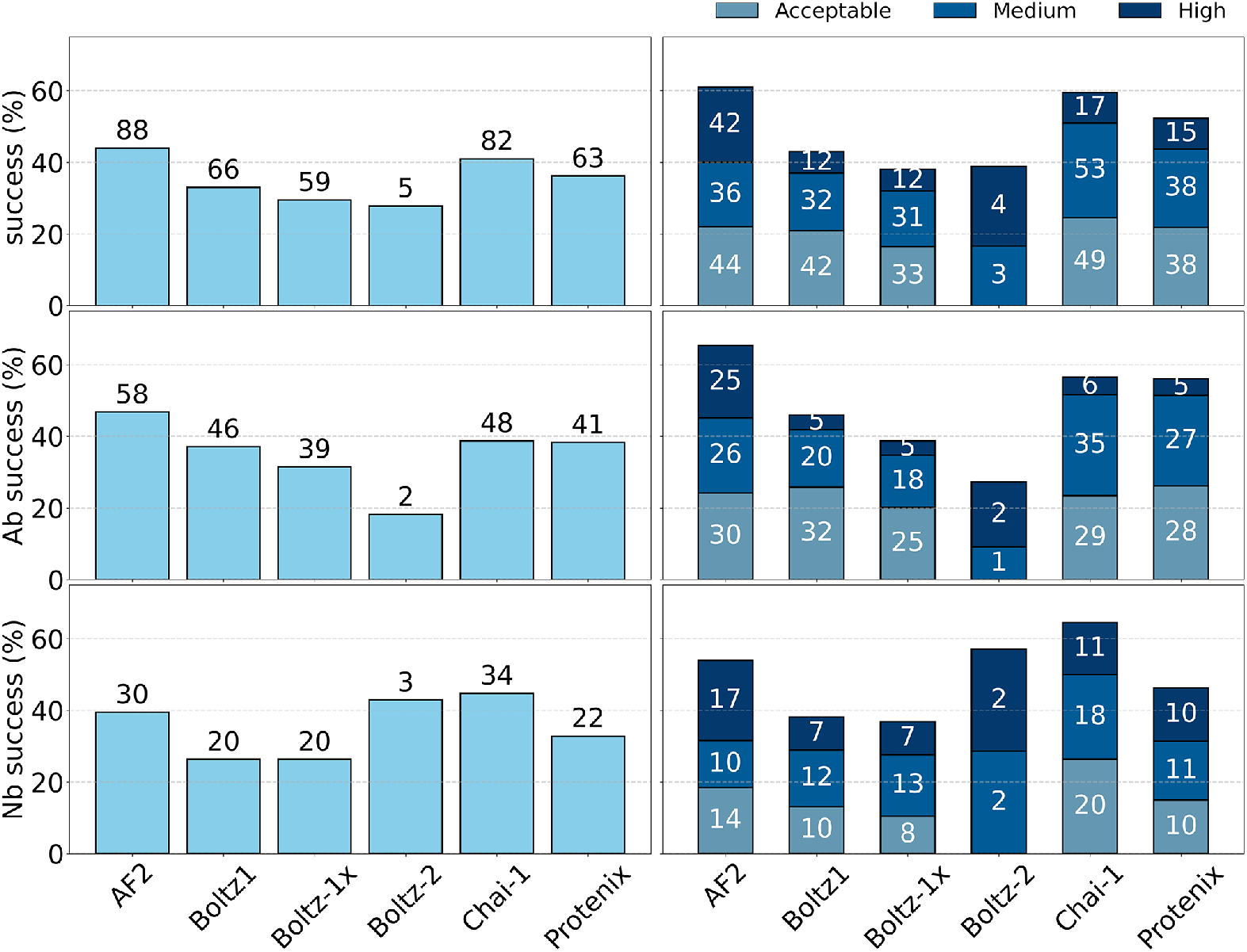
Left panel shows the success rates of all methods across 200 complexes (Boltz-2: 18, Protenix: 174); middle panel shows success rates for antibody complexes (n=124, Boltz-2: 11, Protenix: n=107); right panel displays success rates for nanobody complexes (n=76; Boltz-2: 7, Protenix: n=67). Success rates represent the proportion of unique targets predicted by acceptable, medium, or high accuracy according to CAPRI criteria. Bars were annotated with the target counts.

For all complexes (Fig. 1-top row), AF2 achieved the highest success rate with 88 unique targets. AF2 was followed by Chai-1 (82 targets), Boltz-1 (66), Protenix (63), and Boltz-1x (59). AF2 also produced a significant number of high-quality models (42), demonstrating its ability to capture refined Ab-Ag interfaces. Notably, despite being evaluated on much fewer targets, Boltz-2 also showed a promising performance, yielding only high and medium-quality models. Chai-1 and Protenix were the second and third best methods, respectively, producing high and medium quality models. On the other hand, ESMFold failed to produce any successful predictions for our Ab-Ag complex set, as evaluated by both CAPRI and DockQ criteria (Fig. S1). Although this outcome is not entirely unexpected given that previous benchmarks reporting a poor performance for ESMFold for multimer predictions [39, 41], we repeated ESMFold predictions on a small target subset using an different platform (https://colab.research.google.com/github/sokrypton/ColabFold/blob/main/ESMFold.ipynb) to rule out implementation-related issues. Results from both prediction platforms were identical in terms of complex structures and confidence scores. Therefore, we reported that while ESMFold is highly accurate for monomeric structures [28], it showed no predictive performance for the specific set of Ab–Ag complexes examined here.

AF2 v2.3, an updated version of the model, incorporates pipeline and model modifications, including an increase in training data [42]. Previous benchmarking of AF2 v2.3.0 on a subset of our dataset showed approximately 43% overall success for top-25 models across individual CAPRI levels [29]. In our current benchmarking, however, AF2 v2.3.2 performed much better, achieving over 60% total success across CAPRI levels (Fig. 1-right), which close to the AFsample [43] in the same study [29]. While both versions yielded a comparable 38% success rate for the top-1 tier, AF2 v2.3.2 demonstrated higher success rates for the top-5 tier. These observed differences between the accuracies of two AF2 v2.3 versions could stem from variations in parameters, such as random seed initialization. Additionally, our use of ColabFold, which differs from the original AF2 pipeline in its construction and pairing of MSAs [31], might also contribute to this discrepancy.

We also calculated success rates focusing on specific interaction types, antibody (Ab) and nanobody (Nb) complexes (Fig. 1-middle/bottom rows). For Ab complexes (Fig. 1-middle), AF2 again emerged as the most successful method with a 47% (58/124) success rate. Second best performer was Chai-1 (39%, 48/124), which was closely followed by Protenix (38%, 41/107) and Boltz-1 (37%, 46/124). Notably, Boltz-1x showed a lower performance than Boltz-1, resulting in a total of 39 correctly predicted Ab complexes (31%). Conversely, Boltz-2 showed a weaker performance than Boltz-1 variants, achieving only a 18% success rate (2/11) in the Ab category. When examining the CAPRI-based quality of these predictions, AF2 maintained its performance, yielding again substantial number of high and medium accuracy models. On the other hand, other methods, fall behind AF2’s performance in predicting refined Ab complexes, particularly producing limited number of high quality models.

In the Nb-level assessment (Fig. 1-bottom row), Chai-1 yielded the highest success rate (45%, 34/76), outperforming AF2 (39%, 30/76), while all other methods generally showed lower performance on the Nb complexes compared to their performance on the Ab complexes. Along with this observation, AF2 was still the best method in producing the largest number of high quality models, which is closely followed by Chai-1. We note here that, the performance drop observed for Boltz-1x in the Ab cases was absent for Nb complexes, showing a similar level of success for both Boltz-1 variants. Despite the limited tested complexes, Boltz-2 showed a higher success in Nb category, correctly predicting 3 out 7 targets.

In a recent benchmark of 70 nanobody (Nb) complexes, AF2 v2.3.2 achieved a 33% success rate [44]. In our 76 Nb complexes, we found that AF2 v2.3.2 correctly predicted 30 targets, yielding a 39% success rate (Fig. 1-bottom). Furthermore, AF2 v2.3.2 produced much higher number of high-quality models in our benchmark (24%, 17/76) compared to the 11% in the previous study. These differences in AF2 performance is likely to be stemmed from variations in calculation parameters such as the number of recycling iterations and the fact that the earlier benchmark used DockQ thresholds as its ground truth, whereas we used CAPRI criteria. Noting that particularly models with DockQ scores near boundary values were misclassified in our benchmark (Fig. S1), the use of different ground truths may partly explain the performance difference of AF2 v2.3.2. The same study reported a 47% success rate for AF3, which is slightly below Chai-1’s 49% on our Nb complexes (Fig. 1-bottom). By quality tier, AF3 produced 21% high-quality and 36% medium-quality models [44], while Chai-1 yielded 14% high-quality (11) and 24% medium-quality (18) models. Although our benchmark and the prior study [44] used different ground truths that do not perfectly agree (Fig. S1), we conclude that Chai-1 matches AF3’s performance in Nb complex modeling, and outperforms other AF3-based methods in our Nb complexes.

We visualized the top-scoring predicted model for all methods for a Nb complex (Fig. 2). Consistent with the general success rates reported in Fig. 1, AF2 and Chai-1 achieved highly accurate predictions for this complex. In contrast, Protenix produced an incorrect prediction, despite yielding a marginally incorrect DockQ score. The Boltz variants demonstrated low accuracy for this particular nanobody complex.

**Fig. 2.**
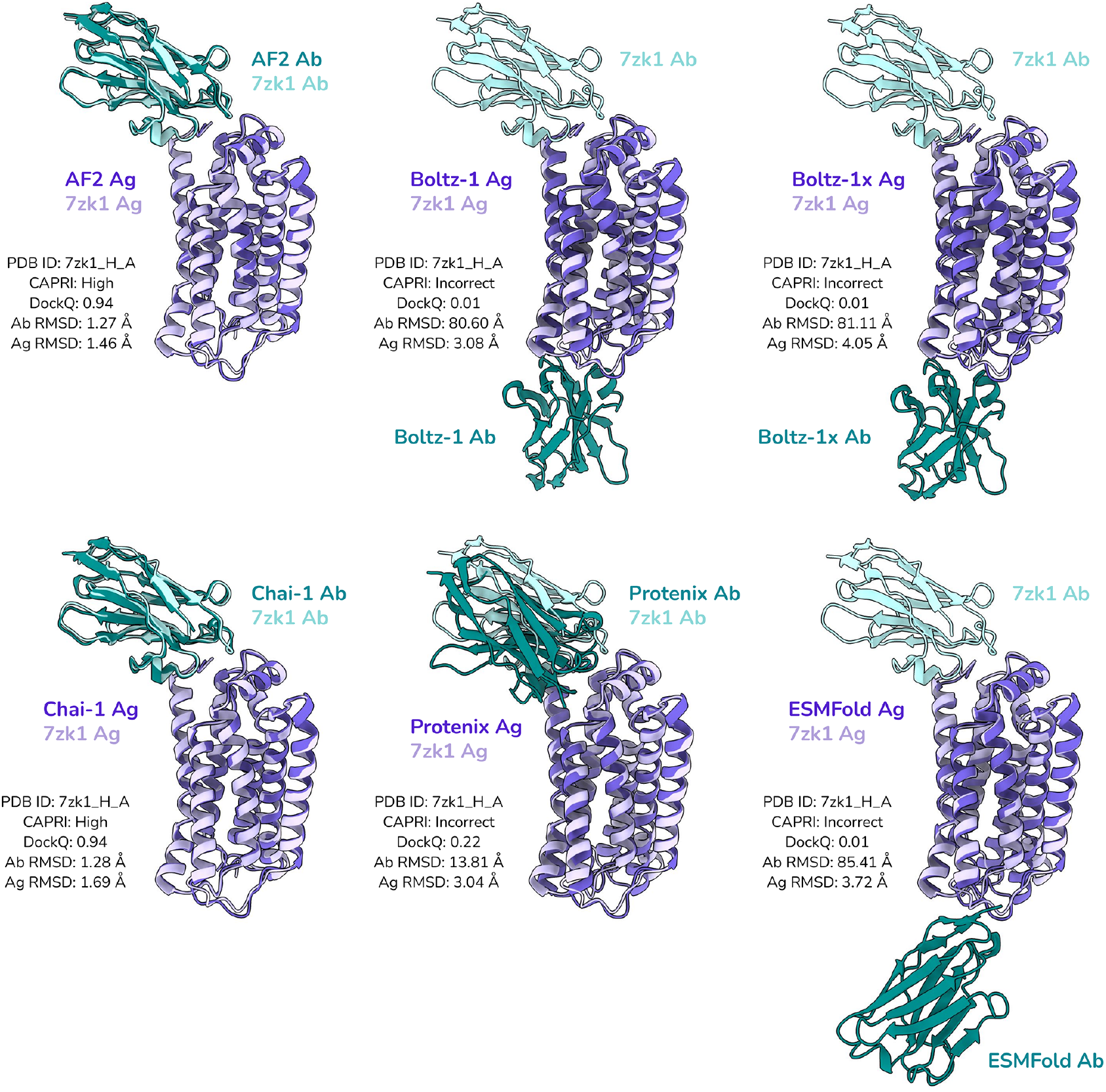
Structural visualization of one complex (PDB ID: 7zk1) showing the top-ranking models. Native structure is depicted in lighter colors, predicted models in darker shades. Ag, antigen; Ab, antibody. C*α* RMSD was given for each subunit.

Taken together, these results demonstrate that AF2 consistently performs well, particularly in generating high-quality Ab-Ag complexes. Chai-1 emerges as a strong competitor, especially in Nb modeling it outperforms all other methods including AF2. Protenix generally showed comparable success in Ab cases with Chai-1, while falling behind it in the Nb interactions. Following these three methods, the Boltz variants show variable performance, with Boltz-1 outperforming Boltz-1x in the Ab category. Despite its being tested in a limited number of complexes, Boltz-2 outperformed earlier Boltz versions in the Nb category.

### Effect of Interface Glycans on Success Rates

In our Ab-Ag complex dataset, we identified 41 interfaces containing glycan residues within 10 Å of the binding interface. Glycosylation, a key post-translational modification (PTM) for antibodies and nanobodies, significantly influences their activity, including their ability to bind target antigens and engage immune effector mechanisms [45]. These PTMs including glycosylation can also impact the accuracy of prediction methods. While AF3-based methods can account for glycosylation during complex modeling, we did not provide any PTM information to the models. Hence, to specifically test how the glycosylated interfaces affect success rates, we conducted an additional analysis to compute success rates for complexes both with and without interface glycans (Fig. S2-S3).

Our analysis revealed a general decline in prediction success for Ab-Ag complexes harboring glycans within 10 Å of the binding interface, relative to those lacking interface glycans (Fig. S2). Protenix stood out as the least affected method, maintaining consistent success rates of 35% (11/31) for glycosylated and 36% (52/143) for non-glycosylated interfaces. Conversely, other methods exhibited reduced success with glycosylated interfaces. Interestingly, AF2 and Chai-1 produced a higher proportion of high-quality models for glycated complexes. In contrast, Boltz-1 variants yielded no high-quality models for glycated interfaces, highlighting their sensitivity to interface PTMs and suggesting that incorporating PTM information could enhance their performance.

To further demonstrate these trends, we examined a specific glycated complex of a nanobody and SARS-CoV-2 RBD (PDB ID: 7z2l) with an N-linked glycosylation at the 434^*th*^ position of RBD (Fig. S3). Without prior glycosylation information, Boltz-1 variants and ESMFold failed to generate correct predictions for this complex. Conversely, AF2, Chai-1, and Protenix successfully produced medium or high-quality models in the absence of explicit glycation input.

### Effect of Depth, Recycling Iterations and Seed

We calculated success rates for the models in top-ranking tiers (Top1 and Top5). Fig. S4 displays the unique target counts across all CAPRI categories, as well as for each individual category. The depth of prediction, meaning the number of predicted models, positively impacted the performance of all methods, resulting in higher success rates. The original AF3 method previously achieved a 60% success rate in antibody-antigen complex prediction using a 1,000-seed sampling strategy [23]. However, a recent benchmark reported a significantly lower success rate of ~ 11% with single-seed sampling [30], underscoring the sensitivity of the method to sampling depth. Our findings similarly indicated that AF2 and all AF3-implementations responded well to an increased number of models in a given prediction (Fig. S4). As such, all methods showed an increase in correctly predicted unique targets as the number of models were increased for a single-seed prediction.

Recycling, which iteratively refines the model by reintroducing the outputs into the network for further optimization [21, 22], appears to be an important factor contributing to the success of Ab-Ag modeling [29]. To assess the impact of recycling on each model’s performance, we compared predictions generated with three recycling iterations to those generated with 20 recycling iterations (Fig. S4). Increasing the number of recycling iterations from 3 to 20 led to an improvement in the performance of all methods, with Chai-1 being the most noticeable. Specifically, Chai-1’s performance increased from 45 to 67 unique targets for the top 1 ranking models and similarly from 50 to 70 for the top 5 and from 70 to 82 for all 25 models (Fig. S4-top). Protenix’s predictions also responded well to increased recycling iterations, leading to increase in the overall target count from 29 to 46 for top 1 tier and from 41 to 57 for the top 5 tier. Not as significant as Chai-1 and Protenix, Boltz-1 also led to improved success upon increased recycling iterations. However, AF2 and Boltz-1x were less responsive to changes in the recycling iterations. As such, AF2 showed slightly improved overall success for 20 recycles, while Boltz-1x showed a slight reduction in success rates particularly for top-1 and top-25 tiers. These results show that although additional recycling generally boosts accuracy in both AF2 and AF3-based methods, it may also impair performance, as illustrated by Boltz-1x’s predictions.

We also evaluated the effect of random seed variability on AF2 performance by performing an additional prediction with 3-recycle iterations (Fig. S4, AF2(3)^*^). While overall success, i.e. unique target count, remained similar in both 3-recycle runs, we noted that distribution of CAPRI quality levels differed upon changing the seeds, suggesting that random seed selection can influence prediction accuracy.

The definition of “success” is critical for accurately evaluating method performance. One way to report it is to count targets for each CAPRI category [29, 30]. While quantifying the number of high/medium quality models within a prediction is valuable for understanding the models’ ability to produce refined models, unique target counts calculated for each CAPRI category carries the risk of overestimating a method’s overall performance. Our analysis suggests that the success rate when reported as the sum of unique target counts in each CAPRI category (Fig. 1-right) inflates the performance compared with success rate computed from unique target counts with at least acceptable accuracy (Fig. 1-left). For example, our benchmark reflected that AF2, when accounting for each CAPRI category individually, correctly predicted 124 targets (Fig. 1-right); however, only 88 of these were unique targets (Fig. 1-left). This inflation occurs because a single target is counted in multiple CAPRI categories. For a more accurate representation of performance, we therefore define the overall success rate as the total number of unique targets predicted with at least acceptable accuracy. This ensures that each success is counted only once, offering a realistic appraisal of a method’s true predictive power especially when multiple models were generated per target.

### Structural Accuracy and Length of CDR-H3 Against Model Quality

CDR-H3 loop is one of the determinants of antibody specificity. Its high sequence and structural diversity make it a central challenge in computational antibody design [6, 16, 20]. To assess the method’s ability to correctly model CDR-H3 loop’s native conformation, we calculated its RMSD relative to the native antibody structure (Fig. 3, top).

**Fig. 3.**
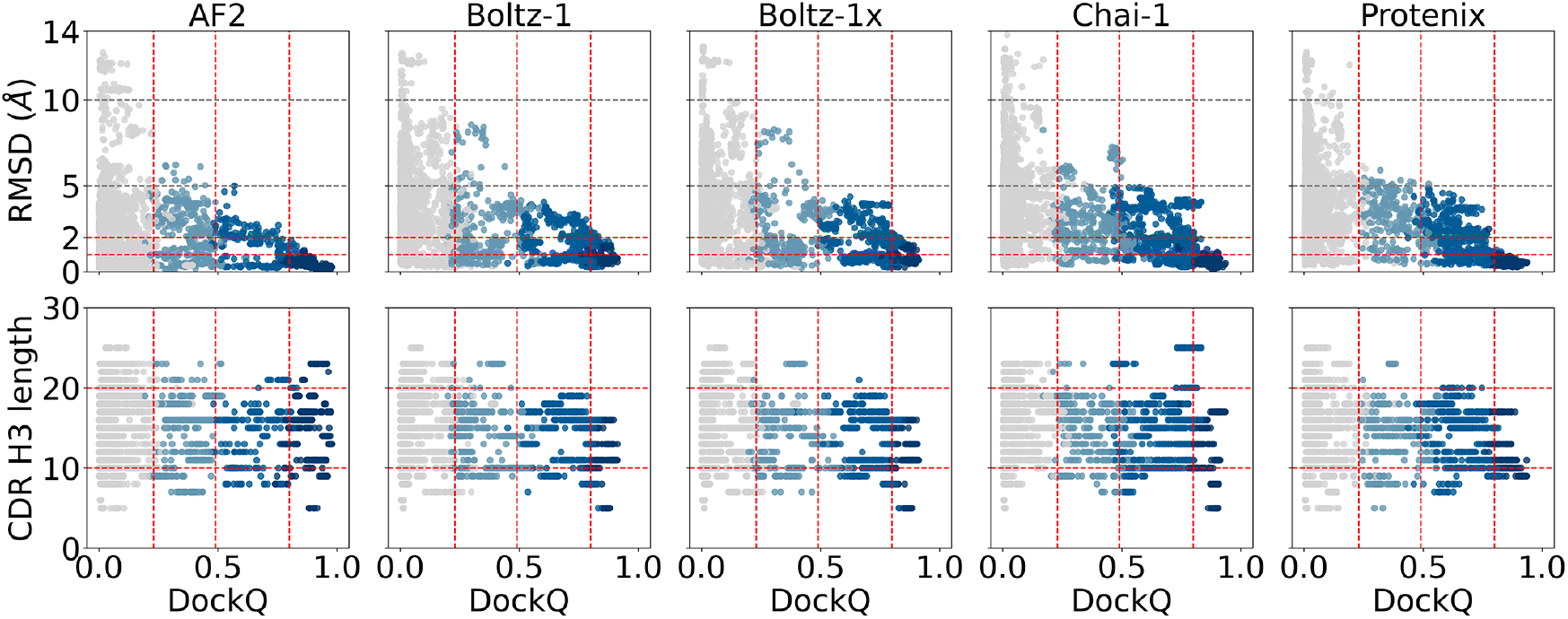
(Top) The structural accuracy of the CDR-H3 loop is plotted against DockQ. Structural accuracy is calculated as the C*α* RMSD of the loop with the native antibody as a reference. Red horizontal lines mark 1 and 2 Å (Bottom) Length of the CDR-H3 loop is plotted against DockQ. In both panels, points are colored according to their CAPRI quality classification, and vertical lines delineate the DockQ thresholds for acceptable (0.23), medium (0.49), and high-quality (0.80) models.

Across all methods, loop RMSD and DockQ score were inversely correlated (Fig. 3, top). For high-quality models, every method captured CDR-H3 conformations within 2 Å of the native conformation. AF2, Chai-1 and Protenix produced a substantial fraction of high-quality models with RMSDs below 1 Å, while Boltz-1 variants yielded a fraction of targets with CDR-H3 loop accuracy in the 1–2 Å range. Medium-quality models from all methods stayed below 5 Å, with AF2 standing out, as almost all medium-quality models of AF2 fell below 3 Å. Nearly all acceptable-quality models from AF2, Chai-1 and Protenix showed RMSDs under 5 Å, whereas Boltz-1 variants exhibited slightly elevated RMSDs in their medium-quality predictions. Finally, incorrect models from every method produced right-skewed RMSD distributions, extending up to 14 Å. One recent study reported a comparable distribution of CDR-H3 RMSD values across different DockQ scores [44]. Another study, which utilized Ab-Ag complexes overlapping with our dataset, evaluated AF3 and found that its CDR-H3 RMSD distributions closely resembled those of AF2 and Chai-1. Therefore, we infer that the CDR-H3 RMSD and DockQ distributions produced by AF3 align with those reported in this work. Overall, this analysis showed that CDR-H3 loop accuracy is strongly correlated with model quality for all methods, reflecting the promise of AF2, Chai-1 and Protenix to find accurate CDR-H3 structures.

We then examined the impact of CDR-H3 loop length on the model quality (Fig. 3, bottom). AF2 and Chai-1 proved robust, consistently generating high and medium-quality models for all loop lengths, short (*<*10 aa), medium (10–20 aa) and long (*>*20 aa). On the other hand, Protenix showed a bias toward medium-length loops (10–20 aa), with few successes for loops shorter than 10 aa or longer than 20 aa. Boltz-1 and Boltz-1x similarly performed best on shorter loops and missed targets whose CDR-H3 length is longer than 20 aa. Notably, only Chai-1 correctly modeled the 25-residue CDR-H3 from the 7mzm structure whilst all other methods failed on this complex. These findings underscore that handling long CDR-H3 loops is a critical differentiator in AI-based Ab–Ag complex modeling and highlights an larger area for improvement for Boltz-1 models.

### Evaluation of Confidence Scores as Predictors of Structural Accuracy

To evaluate how well model confidence scores reflect structural accuracy, we analyzed the relationship between DockQ scores and the confidence measures, namely pTM, ipTM, model confidence, ipLDDT, normalized iPAE and the post hoc score pDockQ2 (Fig. 4). All score distributions except pDockQ2 share one concerning feature for all methods. Incorrect models often receive high scores, which creates a risk of false positives, i.e. low precision. In contrast, pDockQ2 successfully assigns low scores to incorrect models, but at the cost of underrating correct ones. Many high- and medium-quality predictions end up with near-zero pDockQ2 values. This false-negative tendency also appears in iPAE, which scores both high- and medium-quality models poorly. Because pDockQ2 is obtained by fitting a sigmoid curve to the product of ipLDDT and iPAE scores from the antigen chain [39], its failure to reward correct models apparently originates from the iPAE factor (Fig. 4). In other words, pDockQ2’s strength, i.e. high precision via low scores for incorrect models, is undermined by its low recall, as it too often penalizes correct predictions. By contrast, scores such as pTM, ipTM, model confidence, and ipLDDT consistently score correct models highly, with pTM showing the clearest placement of true positives in the high-score range. To balance these opposing risks, which are false positives with pTM/ipTM/model confidence/ipLDDT/iPAE and false negatives with iPAE/pDockQ2, we developed a composite metric called Antibody Confidence (AntiConf). Computed as a weighted average of pTM and pDockQ2, AntiConf captures the precision of pDockQ2 and the recall of pTM, offering a more reliable overall confidence score (Figs. 4).

**Fig. 4.**
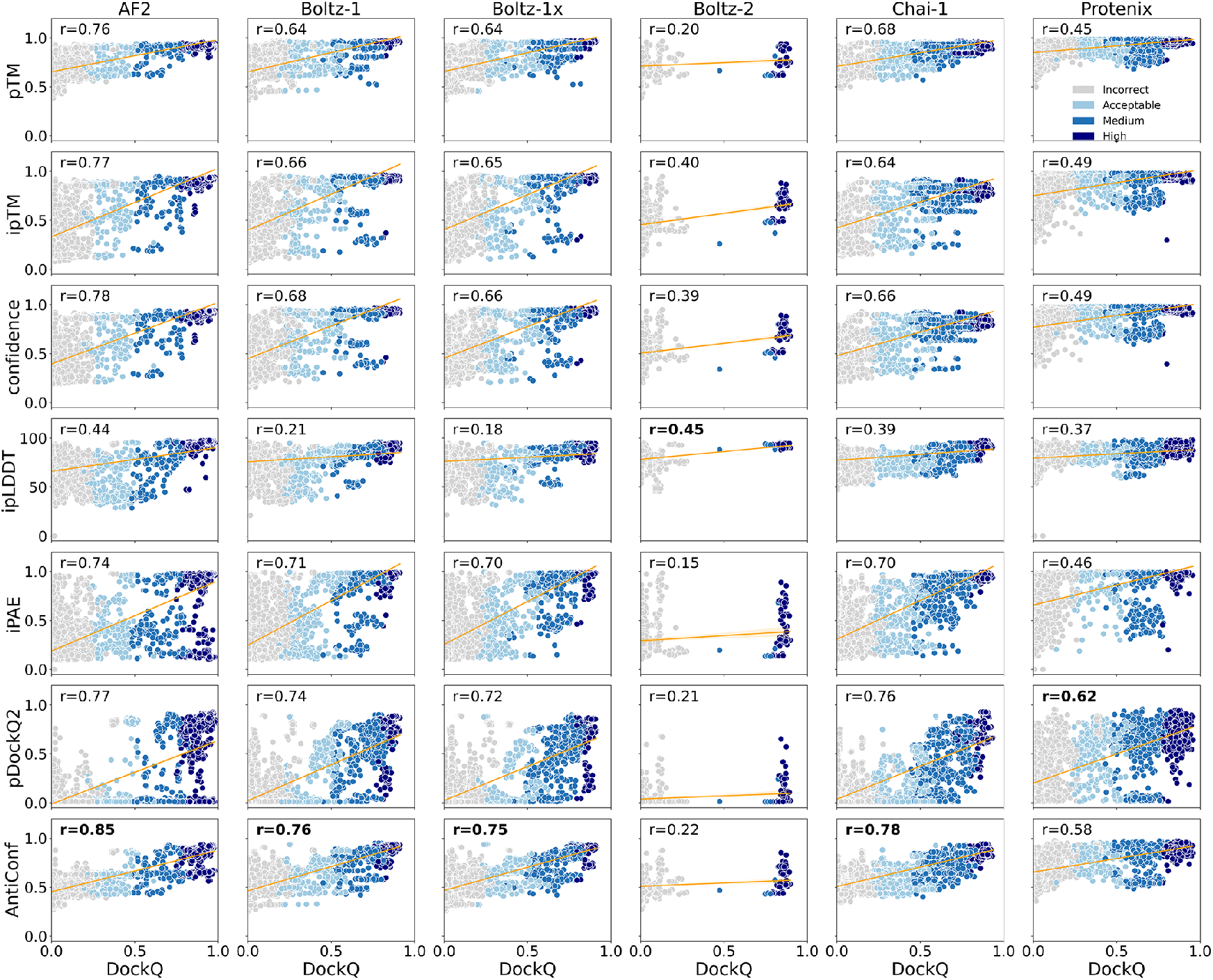
Scatter plots of DockQ scores and confidence metrics across six structure prediction methods. All correlations are significant (*p <* 0.001). ipLDDT, iPAE and pDockQ2 scores for obtained for the antigen chain only.

We reported a consistently stronger associations for the new score AntiConf for all methods except for Protenix (Figs. 4). Notably, for AF2 AntiConf showed a notable increase in the strength of the association. While ipLDDT consistently showed the weakest DockQ association for all methods, except for Boltz-2 which is represented by only a few data points. Following AntiConf, pDockQ2 showed the second strongest correlations with DockQ for all methods (Figs. 4). As we compared the linear correlations for each method, AF2 correlations was consistently the strongest across all scores, which is followed by Chai-1 correlations. Overall, these evaluations implied the power of the composite score, AntiConf, in predicting model quality for these novel antigen-antibody interactions.

The ipTM is a key confidence metric used to evaluate protein–protein interactions by assessing the pTM scores of interface contacts [46]. Because ipTM may include non-interacting flexible regions as part of the interface, its values may not accurately reflect the core interaction region. To address this limitation, alternative scoring approaches considering only the residues directly involved in the interaction have been developed [47, 48]. Among these alternative ipTM scores, we included actifpTM scores for evaluation of AF2 predictions (Fig. S5). ActifpTM showed a weaker association with DockQ compared to ipTM, largely due to a subset of incorrect models with high actifpTM scores.

We also assessed ipLDDT and iPAE scores computed over the entire interface. Notably, the average ipLDDT scores computed over the entire interface correlated more strongly with DockQ compared to antigen-only ipLDDT scores (Fig. S6a), suggesting that averaging ipLDDT over the whole complex yields a more robust indicator of model quality for this Ab-Ag complex set. We additionally investigated iPAE scores computed over the full interface (Fig. S6b). Unlike the trend seen with ipLDDT, antigen-specific iPAE scores were better predictors of model quality than full-interface iPAE scores.

Examining the distributions of model scores across different cycling iterations, we observed that an upward trend in almost all scores for 20-recycle predictions with ipLDDT being the least affected score (Fig. S7). Importantly, we also noted an overestimation of all model scores in the AF3-inspired models compared with AF2, highlighting differences in the internal handling of these metrics by AF3-inspired models. Particularly, scores from Protenix’s models were consistently overestimated compared with the scores from other methods.

### Binary Classification Performance of Confidence Scores

To evaluate how effectively model confidence scores differentiate between correct and incorrect models, we conducted a binary classification analysis based on CAPRI-defined model quality. This allowed us to quantitatively assess the classification performance of the confidence scores (pTM, ipTM, actifpTM, model confidence, ipLDDT, iPAE) and the post-hoc scores (pDockQ2 and AntiConf) (Table S1). Using a fair threshold of 0.80 for all scores, we observed that AntiConf generally yielded the highest precision for all methods, except for Chai-1 and Protenix, for which it was the second-best score, closely trailing pDockQ2. Notably, despite pDockQ2’s higher precision in Chai-1 and Protenix, AntiConf demonstrated a remarkable increase in recall compared to pDockQ2, resulting in an overall high F1 score for both methods. We also computed the proportion of successfully predicted unique targets, termed recall_*target*_, where AntiConf’s performance was also noteworthy. In comparison with the pDockQ2 which gives comparable precision to AntiConf, recall_*target*_ was notably higher in AntiConf, reflecting its potential in recovering higher number of targets without compromising precision (Table S1). Specifically, it returned 39 unique targets (from 88 positive) with a precision of 0.961 for AF2. Similarly, it recovered 43 unique targets out of 82 positives with a precision of 0.972 in Chai-1.

We also acknowledged the classification performance of pTM, ipTM, and model confidence scores, particularly due to their higher recall. However, it is crucial to recognize that in real-world applications, particularly when (costly) experimental structures are not available and are required, high precision (minimizing false positives) is arguably more critical than high recall. AntiConf, integrating the recall strength of pTM and precision from pDockQ2, offers a compelling solution to the reduced recall associated with the highly precise pDockQ2. Therefore, we highlight the power of AntiConf in scenarios where misclassifying incorrect structures as correct could lead to superfluous experimental follow-ups.

Although extensive, this dataset may not reflect the full diversity and complexity of real-world Ab–Ag interactions, potentially underestimating scoring challenges; therefore, our second goal was to evaluate the robustness of AntiConf and other scores under stringent precision conditions. To do this, we identified the lowest threshold for each score and method that resulted in a precision of 1 (Table 1). This specific scenario, guaranteeing 100% precision for each score, would naturally limit recall performance. Thus, as expected, we observed significantly low recall values for every score and method (Table 1). Despite this demanding scenario, AntiConf demonstrated strong predictive value, also maintaining high recall performance compared to other scores. AntiConf either performed the best or the second in terms of recall and recall_*target*_, meaning it identified a large number of unique targets. For example, with AF2, AntiConf yielded the second-highest recall but ultimately identified the highest number unique targets (27), similar to ipLDDT, which showed the highest recall.

**Table 1.** Classification metrics based on the lowest score threshold at precision 1.

| method | score | threshold | recall | F1 | targets | recall<sub>target</sub> |
| --- | --- | --- | --- | --- | --- | --- |
| **AF2**<br>**(88)** | pTM | 0.935 | 0.147 | 0.257 | 16 | 0.182 |
|  | ipTM | 0.916 | 0.168 | 0.288 | 21 | 0.239 |
|  | model conf. | 0.919 | 0.159 | 0.274 | 20 | 0.227 |
|  | iPAE | 0.98 | 0.032 | 0.061 | 4 | 0.045 |
|  | ipLDDT | 0.952 | **0.244** | **0.392** | **27** | **0.307** |
|  | pDockQ2 | 0.848 | 0.152 | 0.264 | 19 | 0.216 |
|  | AntiConf | 0.879 | 0.235 | 0.381 | **27** | **0.307** |
| **Boltz-1**<br>**(66)** | pTM | 0.969 | 0.036 | 0.07 | 3 | 0.045 |
|  | ipTM | 0.96 | 0.02 | 0.039 | 2 | 0.03 |
|  | model conf. | 0.961 | 0.025 | 0.049 | 2 | 0.03 |
|  | iPAE | 0.991 | 0.039 | 0.076 | 3 | 0.045 |
|  | ipLDDT | 0.964 | 0 | 0 | 0 | 0 |
|  | pDockQ2 | 0.899 | 0.001 | 0.002 | 1 | 0.015 |
|  | AntiConf | 0.914 | **0.085** | **0.157** | **4** | **0.061** |
| **Boltz-1x**<br>**(59)** | pTM | 0.968 | 0.051 | 0.097 | **5** | **0.085** |
|  | ipTM | 0.958 | 0.034 | 0.066 | 3 | 0.051 |
|  | model conf. | 0.96 | 0.038 | 0.073 | 4 | 0.068 |
|  | iPAE | 0.992 | 0.039 | 0.075 | 2 | 0.034 |
|  | ipLDDT | 0.957 | 0.001 | 0.002 | 1 | 0.017 |
|  | pDockQ2 | 0.897 | 0.001 | 0.002 | 1 | 0.017 |
|  | AntiConf | 0.913 | **0.082** | **0.152** | 4 | 0.068 |
| **Chai-1**<br>**(82)** | pTM | 0.962 | 0 | 0 | 0 | 0 |
|  | ipTM | 0.919 | 0 | 0 | 0 | 0 |
|  | model conf. | 0.928 | 0 | 0 | 0 | 0 |
|  | iPAE | 0.956 | **0.223** | **0.365** | **21** | **0.256** |
|  | ipLDDT | 0.968 | 0.017 | 0.034 | 1 | 0.012 |
|  | pDockQ2 | 0.753 | 0.141 | 0.247 | 13 | 0.159 |
|  | AntiConf | 0.874 | 0.159 | 0.274 | 13 | 0.159 |
| **Protenix**<br>**(63)** | pTM | 0.985 | **0.056** | **0.105** | **6** | **0.095** |
|  | ipTM | 0.982 | 0 | 0 | 0 | 0 |
|  | model conf. | 0.982 | 0.001 | 0.002 | 1 | 0.016 |
|  | iPAE | 0.996 | 0.005 | 0.009 | 1 | 0.016 |
|  | ipLDDT | 0.985 | 0 | 0 | 0 | 0 |
|  | pDockQ2 | 0.955 | 0.001 | 0.002 | 1 | 0.016 |
|  | AntiConf | 0.976 | 0.001 | 0.002 | 1 | 0.016 |

To assess the robustness of these scores, we further evaluated their precision, recall, and target performance across varying thresholds. We began with the lowest threshold (T) that achieved a precision of 1, which were listed in Table 1, and then progressively decreased it by 0.05, 0.10, and 0.15 (Figs. 5 and S8-S12). As anticipated, the recall performance for all scores monotonically increased with decreasing thresholds. However, a concurrent decrease in precision was observed for all scores, except for pDockQ2 and AntiConf. Both pDockQ2 and AntiConf restored high precision even with a 0.15 decrease in the score threshold. Specifically, for a threshold decrement of 0.15, AntiConf achieved a precision of 0.96 and recalled 42 unique targets for AF2, whereas pDockQ2 recalled 33 targets with a precision of 0.95 (Fig. S8). In contrast, the same condition led to reduced precision for other scores, ranging from 0.59 to 0.86 (Fig. S8).

**Fig. 5.**
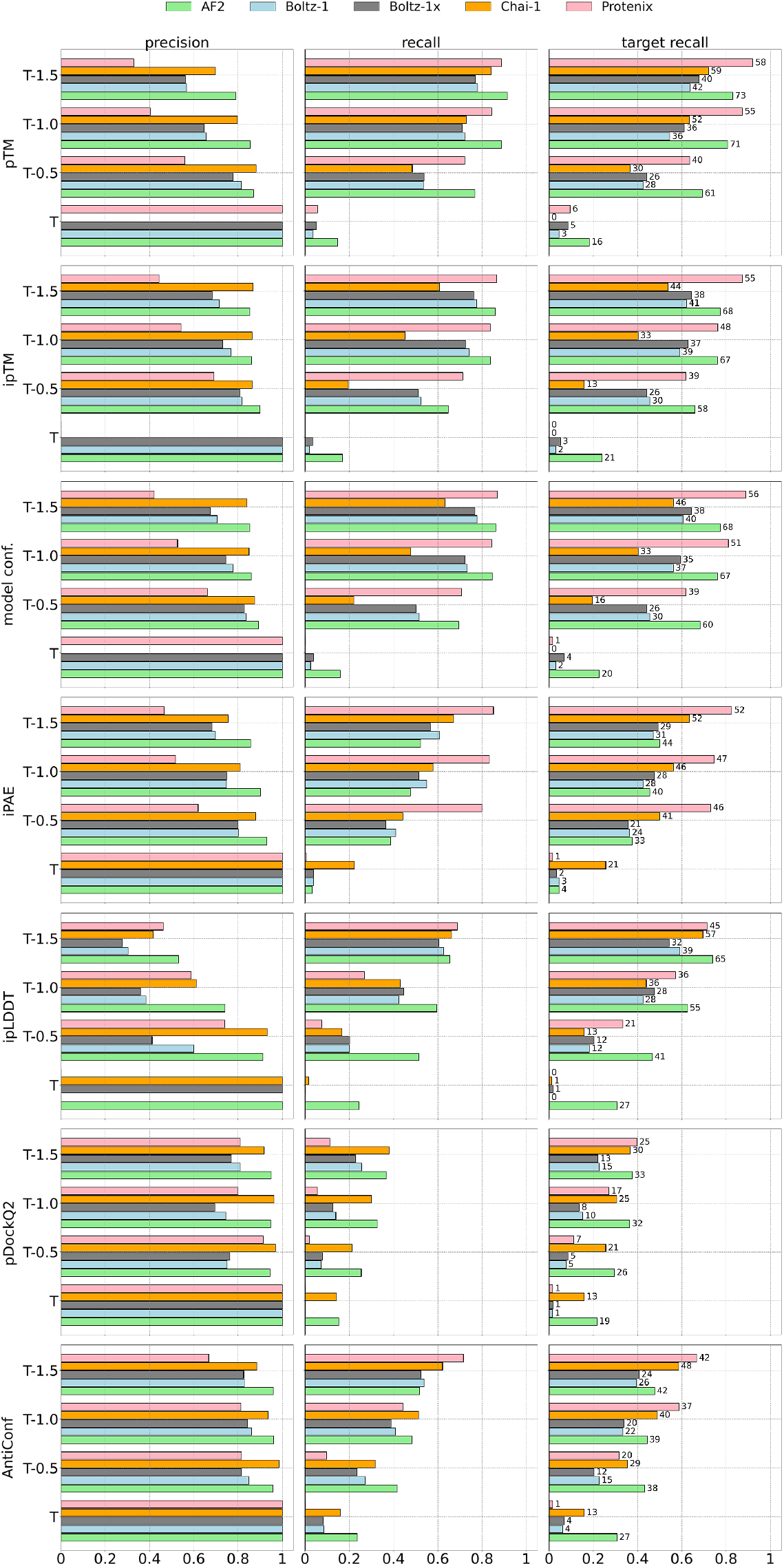
Precision, recall and recall_*target*_ based on different thresholds starting from the lowest threshold that gives precision 1 (T) and decrementing it by 0.05, 0.10 and 0.15. Figs. S8-S12 re-illustrate this plot for each method, with the thresholds explicitly marked.

For AF3-inspired methods, excluding Protenix, AntiConf maintained a stable precision across varying thresholds while improving its recall and recall_*target*_ (Figs. S9-S11). For Chai-1, AntiConf’s precision was slightly lower than that of pDock2 essentially for threshold decremented by 0.10 and 0.15 (Fig. S11). Explicitly, the most divergent result was obtained for T-0.15 threshold that gave a precision of 0.92 for pDockQ2 whilst a 0.88 for AntiConf. Despite this subtle compromise in the precision, AntiConf showed much higher recall and recall_*target*_, i.e. 48 targets compared to pDockQ2’s 30. AntiConf showed the highest precision and recall performance for all thresholds including decremented ones for both Boltz-1 variants (Figs. S9-10). Notably, AntiConf tolerated threshold changes robustly, maintaining a level 0.80 precision across all decremented values. Without compromising precision, it also showed a high recall and recall_*target*_, whilst no other scores were able to maintain a high precision upon threshold decrease.

For Protenix, AntiConf’s robustness against the threshold decrements were less clear as such it showed a larger precision loss for all tested threshold than did pDockQ2 (Fig. S12). Nonetheless, we stressed that AntiConf’s recall advantage was very noticeable against pDockQ2. Even according to the threshold where the largest precision difference was observed (T-0.15), AntiConf yielded a precision of 0.67 against pDockQ2’s 0.81 but it showed a much higher gain in recall and recall_*target*_. Specifically, pDockQ2 predicted 25 correct targets above this threshold whilst AntiConf almost doubled this number with 42 correctly predicted targets. Given the systematic increase in Protenix’s internal metrics compared to other methods (Fig. S7), we underline that a re-calibrated pDockQ2 and AntiConf functions might further ameliorate the performance of these scores for Protenix. Hence, we overall underscore the robustness of AntiConf as pan confidence score that can account for prediction accuracy for Ab-Ag complexes predicted by AF2 and AF3 architectures.

Although evaluated on a small dataset, we present Boltz-2’s score thresholds and classification performance in Table S2. Notably, ipLDDT achieved the highest performance at every threshold, followed by pTM, while other metrics, including AntiConf, showed no particular advantage for Boltz-2 predictions.

We also evaluate another precision-stringent scenario that captures at least 95% precision for all scores (Table S3). If a score could not achieve precision *≥* 0.95, we reported the threshold yielding the second highest precision (Fig. S13). This condition similarly reflected AntiConf’s success for AF2, Boltz-1 and Chai-1 based on recall and recall_*target*_.

Our primary aim in this precision–recall (PR) analysis using progressively lower score thresholds was to assess score robustness and to identify threshold values for future Ab–Ag complex predictions. For the former, PR curves clearly illustrate the trade-off between precision and recall but without mapping back to exact thresholds, they offer limited practical guidance. Moreover, if thresholds are sampled too coarsely, the resulting jumps in the curve can mask narrow regions of peak performance. Thus, we surmise that the detailed precision-recall-recall_*target*_ analysis along with the associated thresholds offer a useful guideline for future Ab-Ag predictions.

We lastly evaluated the precision-recall performance of the confidence scores by concatenating the positive predictions from all methods. We conducted this analysis for the threshold that yield precision of 1 and three incrementally lower thresholds. Using these thresholds, we combined the positives predicted by each method into a single set of positive and obtained performance metrics for each score (Fig. 6). Consistent with the assessments based on individual methods, the collective evaluation revealed that pDockQ2 and AntiConf were the top-performing scores. When predictions from all methods were harmonized, these two metrics delivered the highest recall and recall_*target*_. Both pDockQ2 and AntiConf systematically increased the number of correctly predicted unique targets and the recall rate while maintaining high precision. Notably, AntiConf consistently demonstrated the highest recall_*target*_ among the two across all tested thresholds. This highlights its significant potential for evaluating future antibody-antigen (Ab-Ag) complexes predicted by any of these methods.

**Fig. 6.**
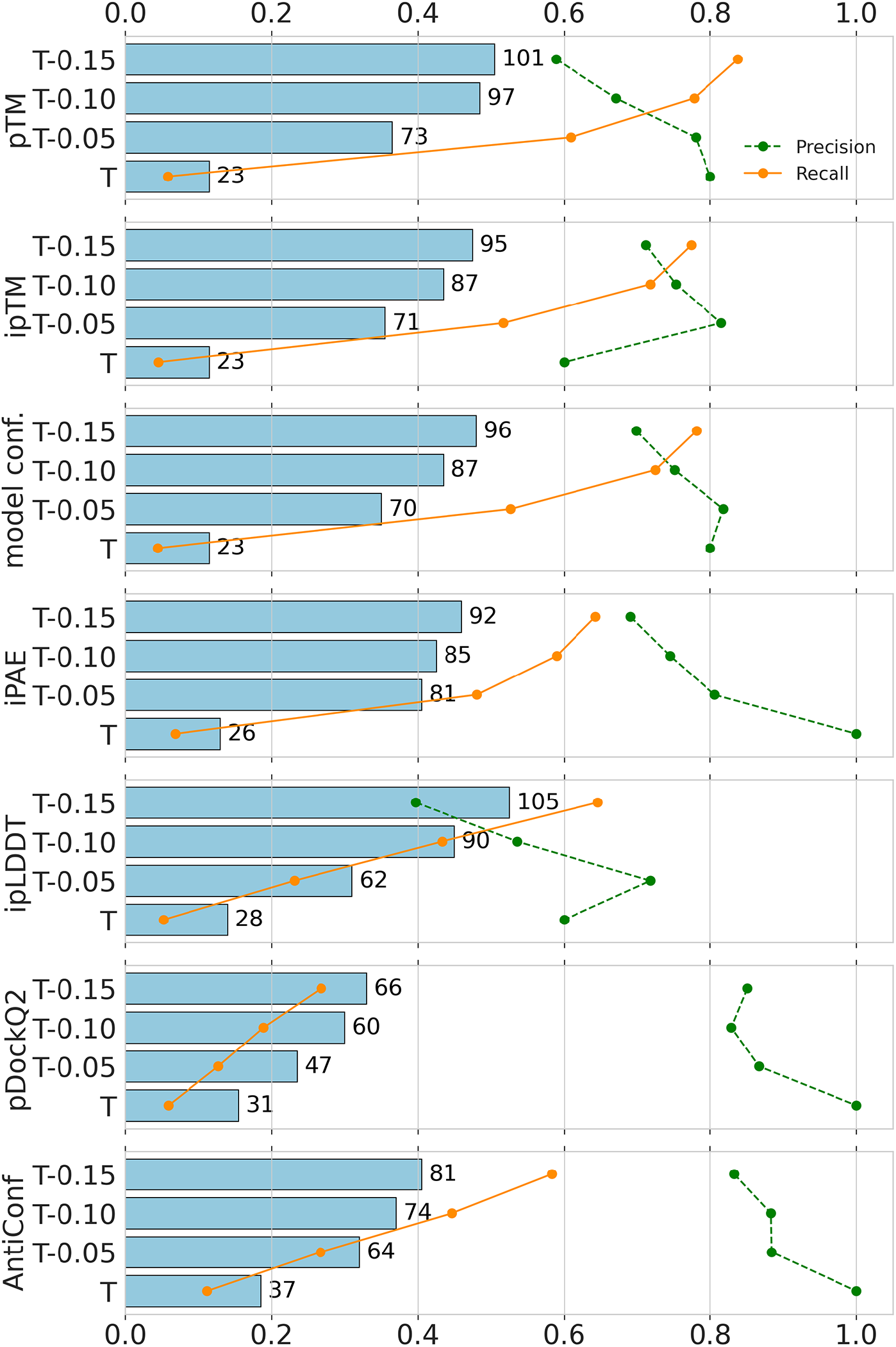
Collective precision, recall and recall_*target*_ performance of all methods based on thresholds starting from the one that gives precision 1 (T) and decrementing it by 0.05, 0.10 and 0.15. See Figs. S8-S12 for corresponding values of T for each method and score.

## Conclusions

Recent advancements in AI-based structure prediction methods, originally designed for single protein structures, have also shown remarkable performance in predicting multimeric protein complexes. However, given that performance gaps still exist, especially for Ab-Ag complex predictions, our study provides a comprehensive comparative assessment of publicly available AF2 and AF3-implementations.

Our study primarily highlights the competitive performance of AF2 and Chai-1 in accurately identifying and refining Ab-Ag complexes. We found that increasing the number of recycling iterations boosts the performance of Chai-1, Protenix, and AF2, while Boltz-1 variants responded differently. Among various model confidence scores from interface-specific to template modeling scores, pDockQ2 demonstrated high precision potential, while pTM offered strong recall. By combining these, we developed a novel score called AntiConf, which stands out as the optimal metric for maximizing recall in challenging scenarios that demand high precision.

AntiConf enables the transformation of sequence data into high-quality structural models, facilitating robust selection of epitope-specific binders. Thus, it can also be used as an effective filtering strategy for antibody design candidates generated by generative models. These capabilities make AntiConf valuable for both computational prediction and experimental validation workflows. Thus, we appraise that AntiConf holds significant promise for accurate determination of correct Ab-Ag complexes from predictions generated by AF2 and AF3-implementations.

While our work points out some limitations in AI-based Ab-Ag complex predictions particularly for Boltz-1 variants, it also demonstrates how current methodologies can be harmonized to maximize prediction accuracy. Combining different prediction methods, rather than relying on a single methodology, it is possible to extract a greater number of correct predictions.

We lastly underscore the importance of future, preferably large-scale, assessment studies to fully understand the generalizability of the confidence scores including AntiConf for estimating the quality of Ab-Ag complex predictions. Nonetheless, the thresholds reported for various confidence metrics in this study offer valuable guidelines for optimizing the performance of different methods for Ab-Ag complexes.

