## Supplementary information for "Confidence Scoring for AI-Predicted Antibody–Antigen Complexes: AntiConf as a Precision-driven Metric"

**Table S1.** Binary classification metrics based on CAPRI incorrect vs rest

| method | score | precision | recall | F1 | NPV | AP | targets | recall$_{target}$ |
|---|---|---|---|---|---|---|---|---|
| **AF2 (88)** | pTM | 0.81 | 0.909 | 0.857 | 0.969 | 0.888 | 73 | 0.83 |
| | ipTM | 0.856 | 0.849 | 0.852 | 0.951 | 0.893 | 67 | 0.761 |
| | actifpTM | 0.332 | 0.941 | 0.491 | 0.952 | 0.469 | 82 | 0.932 |
| | model conf | 0.854 | 0.858 | 0.856 | 0.954 | 0.896 | 68 | 0.773 |
| | ipLDDT | 0.523 | 0.656 | 0.582 | 0.878 | 0.705 | 65 | 0.739 |
| | iPAE | 0.836 | 0.534 | 0.652 | 0.864 | 0.765 | 45 | 0.511 |
| | pDockQ2 | 0.945 | 0.25 | 0.396 | 0.803 | 0.772 | 26 | 0.295 |
| | AntiConf | 0.961 | 0.458 | 0.62 | 0.849 | 0.891 | 39 | 0.443 |
| **Boltz-1 (66)** | pTM | 0.545 | 0.825 | 0.656 | 0.947 | 0.749 | 42 | 0.636 |
| | ipTM | 0.709 | 0.787 | 0.746 | 0.942 | 0.756 | 41 | 0.621 |
| | model conf | 0.694 | 0.788 | 0.738 | 0.942 | 0.762 | 41 | 0.621 |
| | ipLDDT | 0.284 | 0.664 | 0.398 | 0.863 | 0.368 | 41 | 0.621 |
| | iPAE | 0.671 | 0.641 | 0.656 | 0.907 | 0.673 | 35 | 0.53 |
| | pDockQ2 | 0.749 | 0.137 | 0.232 | 0.813 | 0.649 | 10 | 0.152 |
| | AntiConf | 0.865 | 0.453 | 0.595 | 0.872 | 0.751 | 23 | 0.348 |
| **Boltz-1x (59)** | pTM | 0.553 | 0.811 | 0.658 | 0.945 | 0.738 | 42 | 0.712 |
| | ipTM | 0.678 | 0.77 | 0.721 | 0.939 | 0.735 | 38 | 0.644 |
| | model conf | 0.664 | 0.772 | 0.714 | 0.939 | 0.743 | 38 | 0.644 |
| | ipLDDT | 0.262 | 0.614 | 0.368 | 0.848 | 0.342 | 33 | 0.559 |
| | iPAE | 0.634 | 0.597 | 0.615 | 0.898 | 0.652 | 32 | 0.542 |
| | pDockQ2 | 0.726 | 0.124 | 0.212 | 0.814 | 0.626 | 8 | 0.136 |
| | AntiConf | 0.853 | 0.432 | 0.573 | 0.87 | 0.734 | 22 | 0.373 |
| **Boltz-2 (5)** | pTM | 0.31 | 0.355 | 0.331 | 0.865 | 0.302 | 2 | 0.4 |
| | ipTM | 0.2 | 0.0523 | 0.083 | 0.833 | 0.331 | 1 | 0.2 |
| | model conf | 0.323 | 0.132 | 0.187 | 0.842 | 0.333 | 1 | 0.2 |
| | ipLDDT | 0.274 | 1 | 0.431 | 1 | 0.744 | 5 | 1 |
| | iPAE | 0.105 | 0.026 | 0.042 | 0.828 | 0.219 | 1 | 0.2 |
| | pDockQ2 | 0 | 0 | 0 | 0.831 | 0.284 | 0 | 0 |
| | AntiConf | 0.4 | 0.026 | 0.049 | 0.834 | 0.28 | 1 | 0.2 |
| **Chai-1 (82)** | pTM | 0.661 | 0.854 | 0.745 | 0.932 | 0.782 | 63 | 0.768 |
| | ipTM | 0.856 | 0.506 | 0.636 | 0.827 | 0.755 | 36 | 0.439 |
| | model conf | 0.862 | 0.558 | 0.678 | 0.842 | 0.761 | 41 | 0.5 |
| | ipLDDT | 0.385 | 0.732 | 0.505 | 0.827 | 0.612 | 64 | 0.78 |
| | iPAE | 0.752 | 0.683 | 0.716 | 0.875 | 0.818 | 52 | 0.634 |
| | pDockQ2 | 1 | 0.0841 | 0.155 | 0.728 | 0.805 | 9 | 0.11 |
| | AntiConf | 0.972 | 0.447 | 0.612 | 0.815 | 0.855 | 37 | 0.451 |
| **Protenix (63)** | pTM | 0.298 | 0.923 | 0.451 | 0.911 | 0.662 | 61 | 0.968 |
| | ipTM | 0.396 | 0.874 | 0.545 | 0.928 | 0.639 | 56 | 0.889 |
| | model conf | 0.375 | 0.876 | 0.526 | 0.924 | 0.638 | 56 | 0.889 |
| | ipLDDT | 0.348 | 0.817 | 0.488 | 0.886 | 0.512 | 51 | 0.81 |
| | iPAE | 0.429 | 0.87 | 0.575 | 0.933 | 0.66 | 53 | 0.841 |
| | pDockQ2 | 0.817 | 0.122 | 0.212 | 0.769 | 0.699 | 26 | 0.413 |
| | AntiConf | 0.611 | 0.784 | 0.687 | 0.919 | 0.698 | 43 | 0.683 |

Score threshold=0.8; NPV: negative predictive value; AP: average precision.

**Table S2.** Boltz-2 classification metrics based on the thresholds at precision 1

| score | threshold | precision | recall | F1 | targets | $\text{recall}^*_{target}$ |
|---|---|---|---|---|---|---|
| pTM | 0.965 | 0 | 0 | 0 | 0 | 0 |
| | 0.915 | 0.429 | 0.079 | 0.133 | 1 | 0.2 |
| | 0.865 | 0.52 | 0.342 | 0.413 | 2 | 0.4 |
| | 0.815 | 0.338 | 0.342 | 0.34 | 2 | 0.4 |
| ipTM | 0.946 | 0 | 0 | 0 | 0 | 0 |
| | 0.896 | 0 | 0 | 0 | 0 | 0 |
| | 0.846 | 0.222 | 0.026 | 0.047 | 1 | 0.2 |
| | 0.796 | 0.2 | 0.053 | 0.083 | 1 | 0.2 |
| model conf | 0.95 | 0 | 0 | 0 | 0 | 0 |
| | 0.9 | 0 | 0 | 0 | 0 | 0 |
| | 0.85 | 0.182 | 0.026 | 0.046 | 1 | 0.2 |
| | 0.8 | 0.323 | 0.132 | 0.187 | 1 | 0.2 |
| iPAE | 0.984 | 0 | 0 | 0 | 0 | 0 |
| | 0.934 | 0 | 0 | 0 | 0 | 0 |
| | 0.884 | 0.125 | 0.013 | 0.024 | 1 | 0.2 |
| | 0.834 | 0.118 | 0.026 | 0.043 | 1 | 0.2 |
| ipLDDT | 0.955 | 1 | 0.013 | 0.026 | 1 | 0.2 |
| | 0.905 | 0.647 | 0.724 | 0.683 | 3 | 0.6 |
| | 0.855 | 0.429 | 1 | 0.601 | 5 | 1 |
| | 0.805 | 0.287 | 1 | 0.446 | 5 | 1 |
| pDockQ2 | 0.676 | 0 | 0 | 0 | 0 | 0 |
| | 0.626 | 0.333 | 0.013 | 0.025 | 1 | 0.2 |
| | 0.576 | 0.25 | 0.013 | 0.025 | 1 | 0.2 |
| | 0.526 | 0.4 | 0.026 | 0.049 | 1 | 0.2 |
| AntiConf | 0.877 | 0 | 0 | 0 | 0 | 0 |
| | 0.827 | 0.4 | 0.026 | 0.049 | 1 | 0.2 |
| | 0.777 | 0.286 | 0.026 | 0.048 | 1 | 0.2 |
| | 0.727 | 0.357 | 0.066 | 0.111 | 1 | 0.2 |

*Total number correctly predicted unique targets 5.

**Table S3.** Classification metrics based on the lowest score threshold at precision ≥0.95

| method | score | threshold | precision* | recall | F1 | target | $\textbf{recall}_{target}$ |
|---|---|---|---|---|---|---|---|
| **AF2 (88)** | pTM | 0.917 | 0.95 | 0.407 | 0.586 | 41 | 0.466 |
| | ipTM | 0.885 | 0.95 | 0.445 | 0.608 | **47** | **0.534** |
| | model conf. | 0.892 | 0.95 | 0.442 | 0.607 | 44 | 0.5 |
| | ipLDDT | 0.931 | 0.95 | 0.397 | 0.561 | 37 | 0.42 |
| | iPAE | 0.979 | 0.95 | 0.032 | 0.061 | 4 | 0.045 |
| | pDockQ2 | 0.549 | 0.95 | 0.448 | 0.611 | 39 | 0.443 |
| | AntiConf | 0.69 | 0.95 | **0.533** | **0.683** | 45 | 0.511 |
| **Boltz-1 (66)** | pTM | 0.967 | 0.95 | 0.056 | 0.105 | 4 | 0.061 |
| | ipTM | 0.959 | 0.95 | 0.02 | 0.041 | 2 | 0.03 |
| | model conf. | 0.96 | 0.95 | 0.04 | 0.079 | 2 | 0.03 |
| | ipLDDT | 0.964 | 1 | 0 | 0 | 0 | 0 |
| | iPAE | 0.991 | 0.95 | 0.039 | 0.075 | 3 | 0.045 |
| | pDockQ2 | 0.899 | 1 | 0.001 | 0.002 | 1 | 0.015 |
| | AntiConf | 0.912 | 0.95 | **0.093** | **0.169** | **5** | **0.076** |
| **Boltz-1x (59)** | pTM | 0.967 | 0.95 | 0.062 | 0.117 | **6** | **0.102** |
| | ipTM | 0.958 | 0.95 | 0.04 | 0.077 | 4 | 0.068 |
| | model conf. | 0.96 | 0.95 | 0.041 | 0.08 | 4 | 0.068 |
| | ipLDDT | 0.957 | 1 | 0.001 | 0.002 | 1 | 0.017 |
| | iPAE | 0.991 | 0.95 | 0.04 | 0.077 | 2 | 0.034 |
| | pDockQ2 | 0.897 | 1 | 0.001 | 0.002 | 1 | 0.017 |
| | AntiConf | 0.911 | 0.95 | **0.091** | **0.166** | 5 | 0.085 |
| **Chai-1 (82)** | pTM | 0.962 | 1 | 0 | 0 | 0 | 0 |
| | ipTM | 0.919 | 1 | 0 | 0 | 0 | 0 |
| | model conf. | 0.928 | 1 | 0 | 0 | 0 | 0 |
| | ipLDDT | 0.924 | 0.95 | 0.146 | 0.255 | 11 | 0.134 |
| | iPAE | 0.942 | 0.95 | 0.315 | 0.474 | 27 | 0.329 |
| | pDockQ2 | 0.63 | 0.95 | 0.321 | 0.482 | 27 | 0.329 |
| | AntiConf | 0.793 | 0.95 | **0.466** | **0.626** | **39** | **0.476** |
| **Protenix (63)** | pTM | 0.985 | 0.95 | **0.116** | **0.208** | **9** | **0.143** |
| | ipTM | 0.982 | 1 | 0 | 0 | 0 | 0 |
| | model conf. | 0.982 | 1 | 0.001 | 0.002 | 1 | 0.016 |
| | ipLDDT | 0.985 | 1 | 0 | 0 | 0 | 0 |
| | iPAE | 0.997 | 1 | 0.002 | 0.005 | 1 | 0.016 |
| | pDockQ2 | 0.95 | 1 | 0.001 | 0.002 | 1 | 0.016 |
| | AntiConf | 0.976 | 1 | 0.001 | 0.002 | 1 | 0.016 |

*We identified the score thresholds corresponding to a precision of 0.95. If a score could not achieve precision ≥0.95, we reported the threshold yielding the second highest precision.

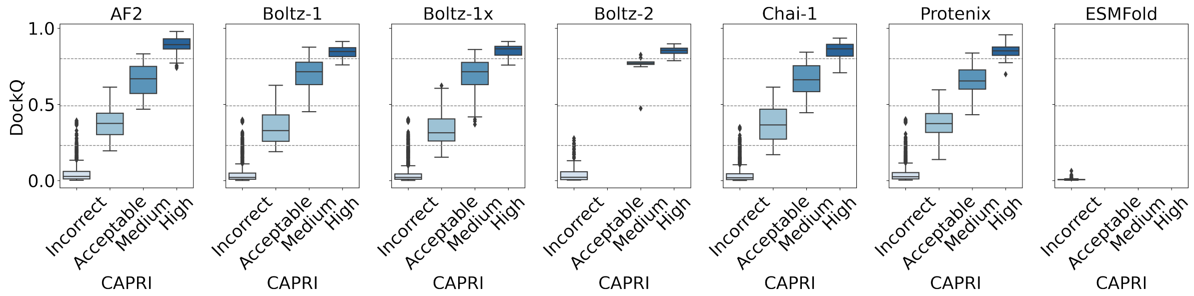

**Fig S1.** DockQ score distributions across CAPRI classes. Dashed lines represent DockQ thresholds (0.23, 0.49, and 0.80).

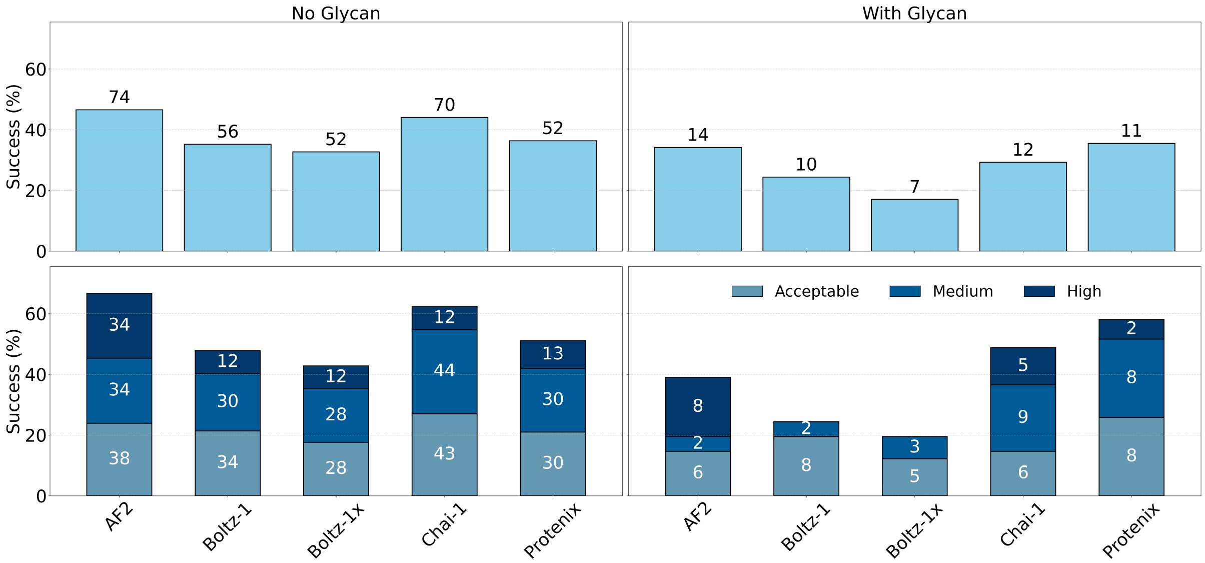

**Fig S2.** Success rates for the complexes without and with interface glycans within 10 Å of the interface.

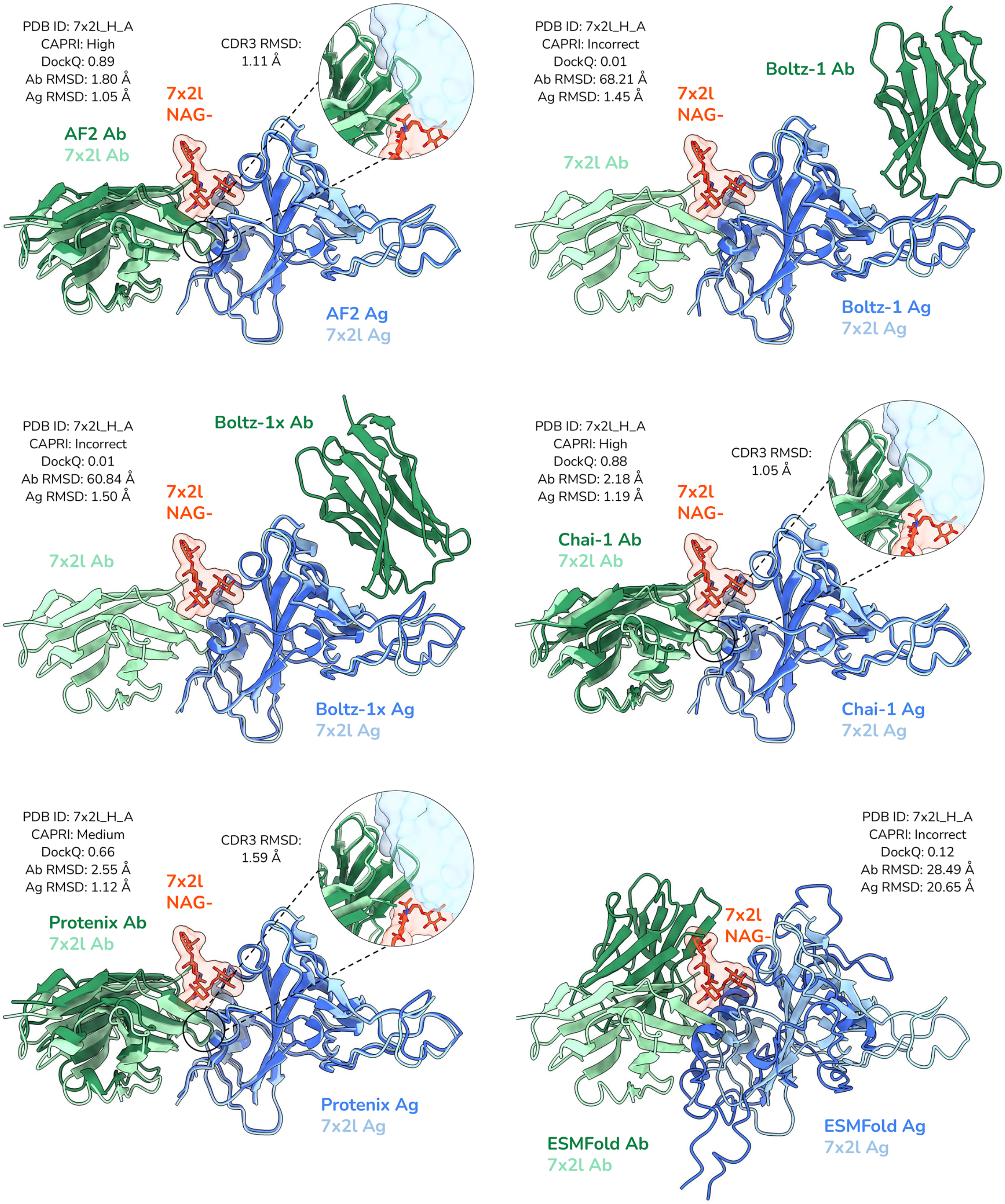

**Fig S3.** An example view of predicted models in comparison with the PDB complex with a glycated interface.

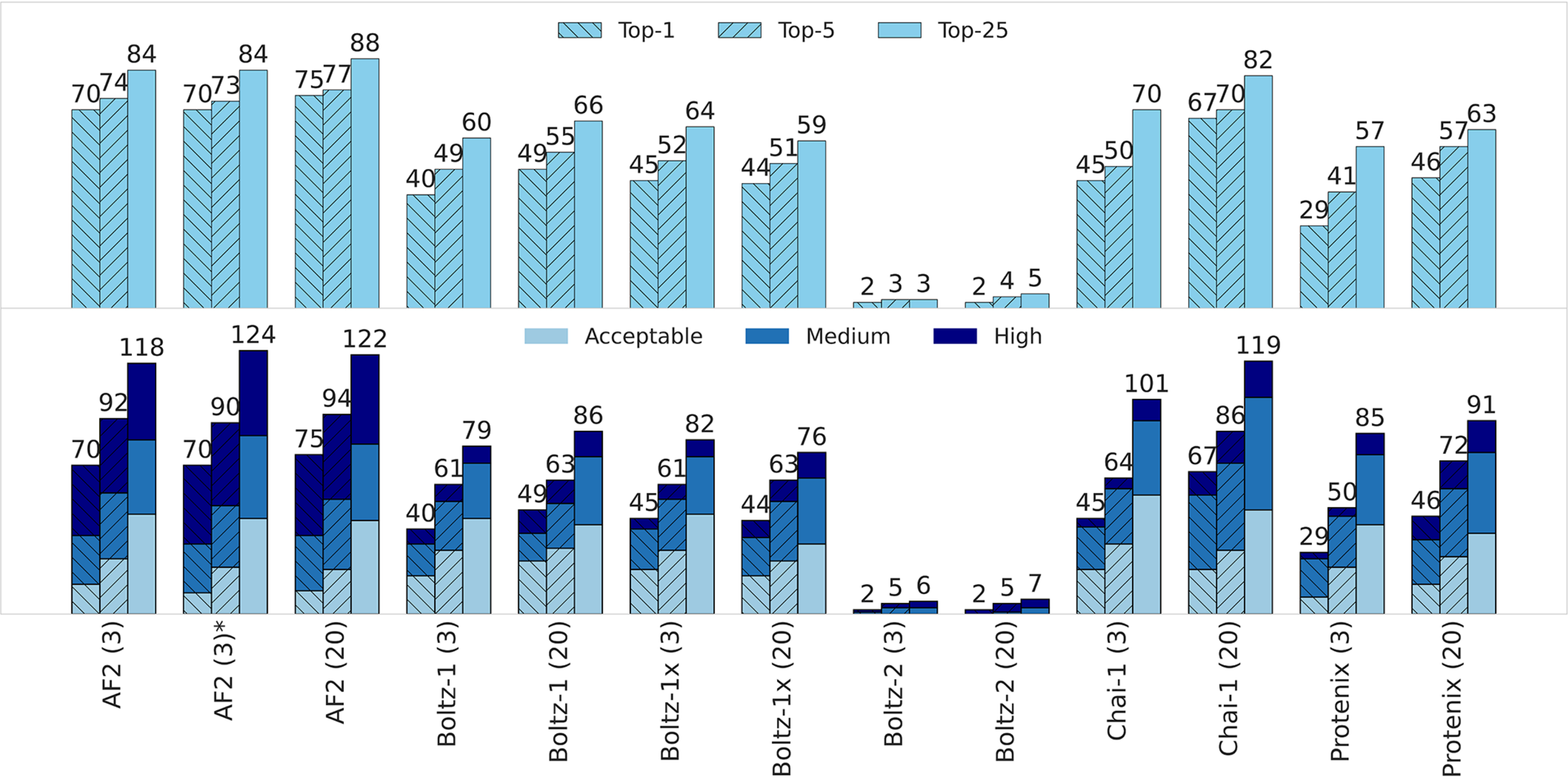

**Fig S4.** Success rates across the top-1, top-5 or top-25 ranked models. Rates are calculated based on the unique targets that achieved (left) at least acceptable level accuracy and (right) a specified accuracy level. Model rankings are based on the model confidence score [?]. Bars are color-coded by CAPRI category, with annotations indicating the number of unique targets.

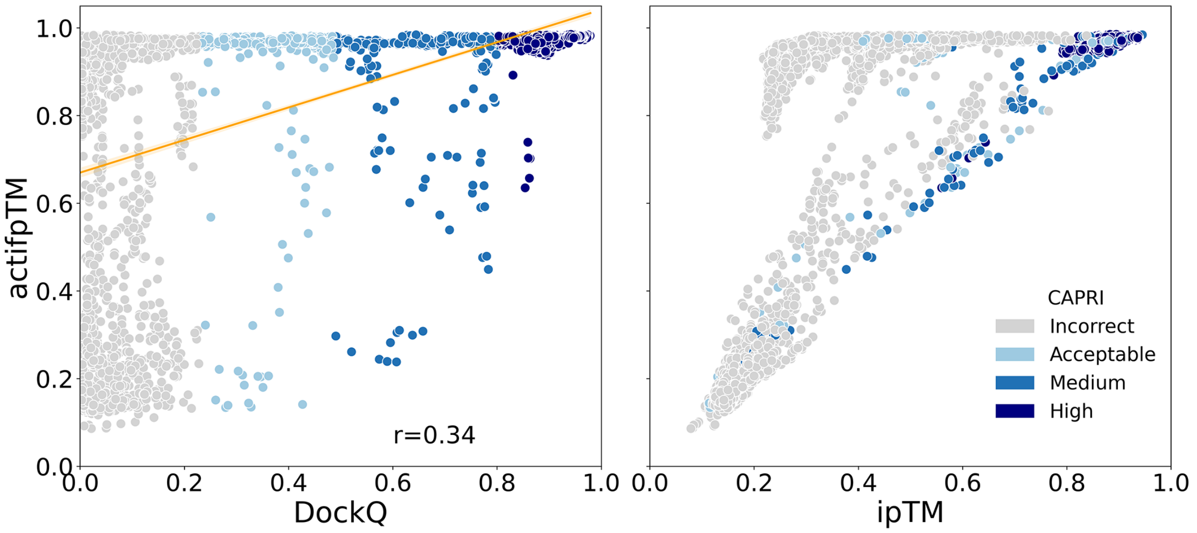

**Fig S5.** Distribution of actifpTM scores versus DockQ and versus ipTM. Scores were given for AF2 predictions with 20 recycle iterations.

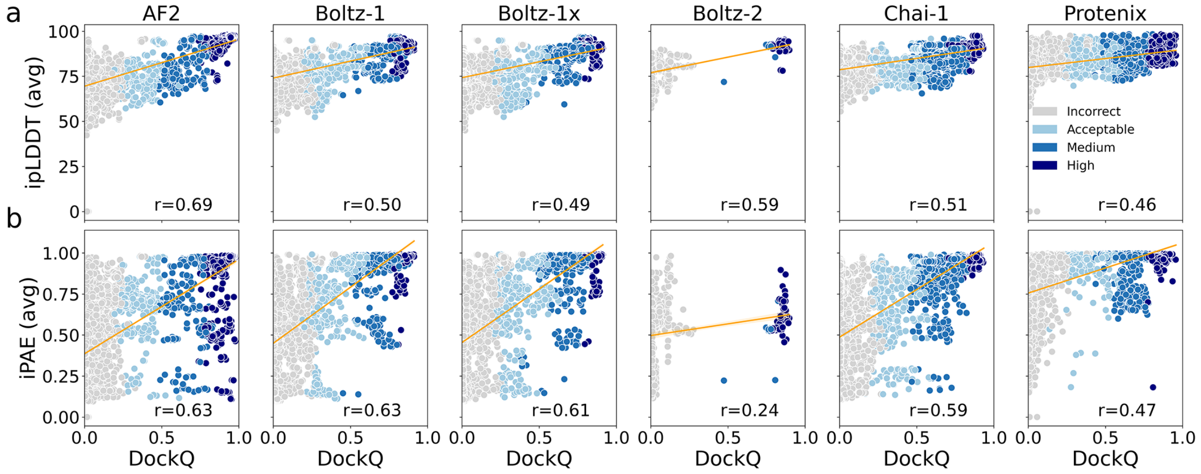

**Fig S6.** Scatter plots of DockQ scores and interface metrics calculated over the entire interface, (a) ipLDDT and (b) iPAE. All correlations are significant ($p < 0.001$).

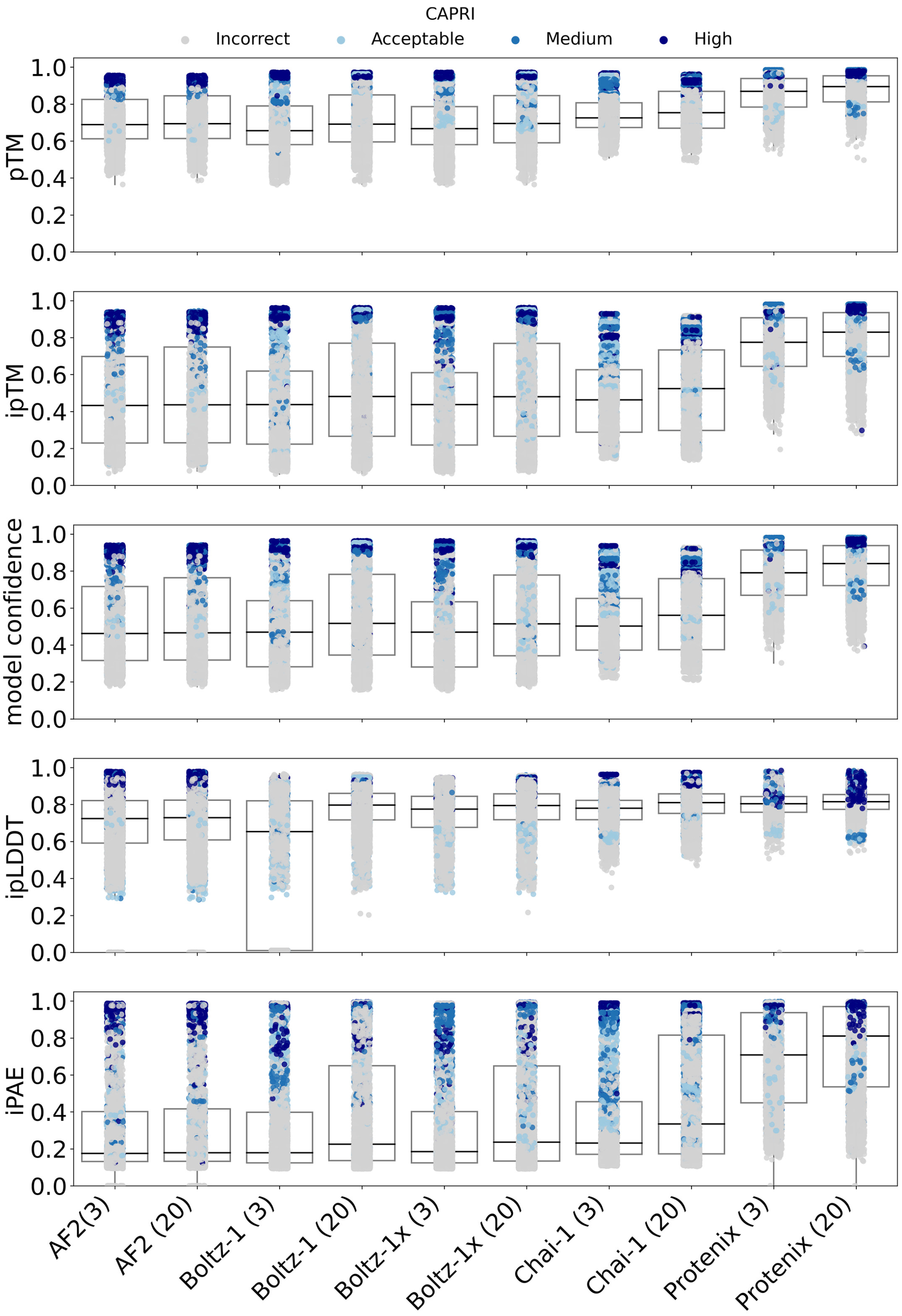

**Fig S7.** Score distributions in all tested predictions.

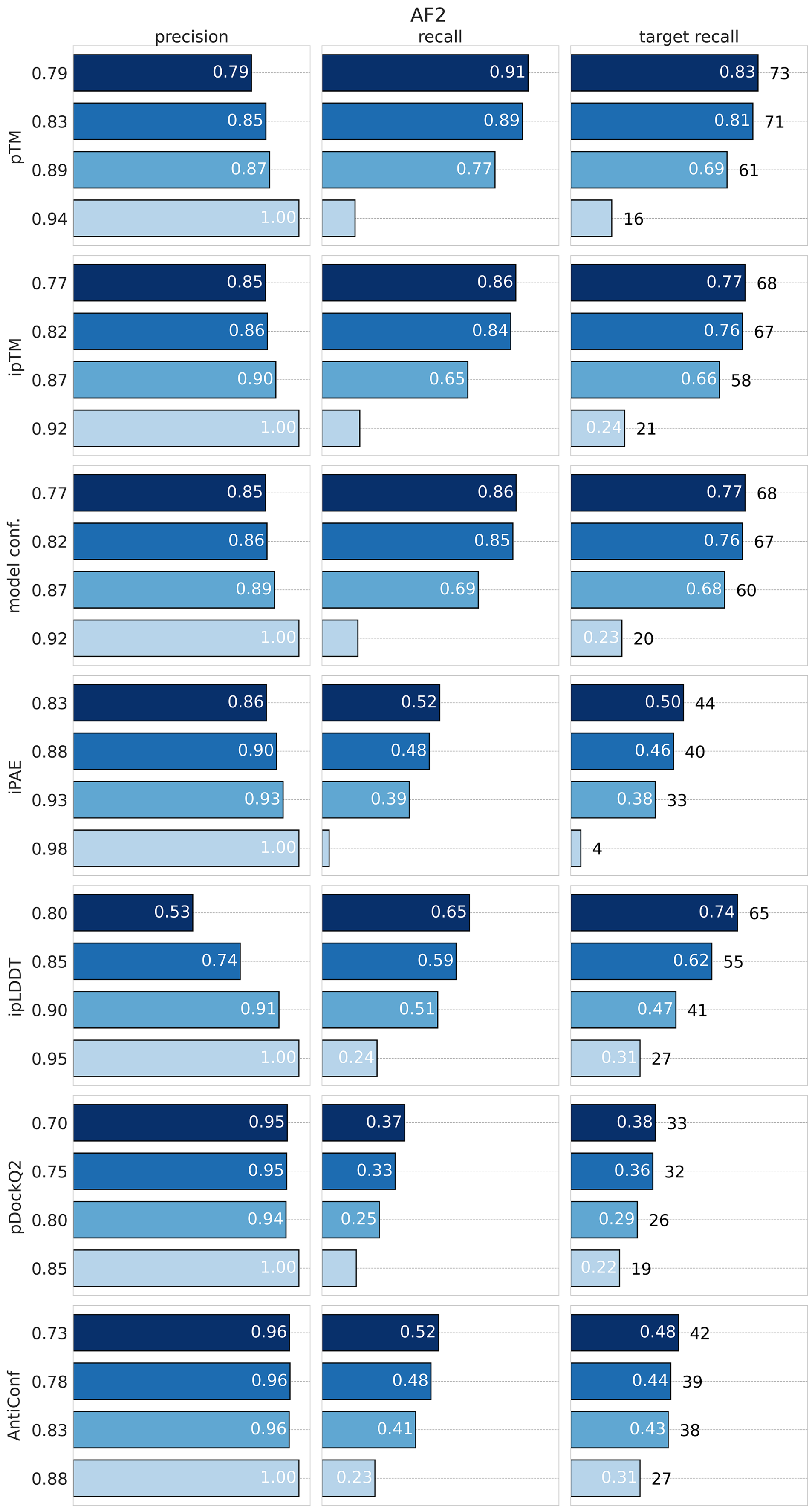

**Fig S8.** AF2 performance using thresholds starting from the lowest value that achieves a precision of 1, and then decrementing it by 0.05, 0.10, and 0.15.

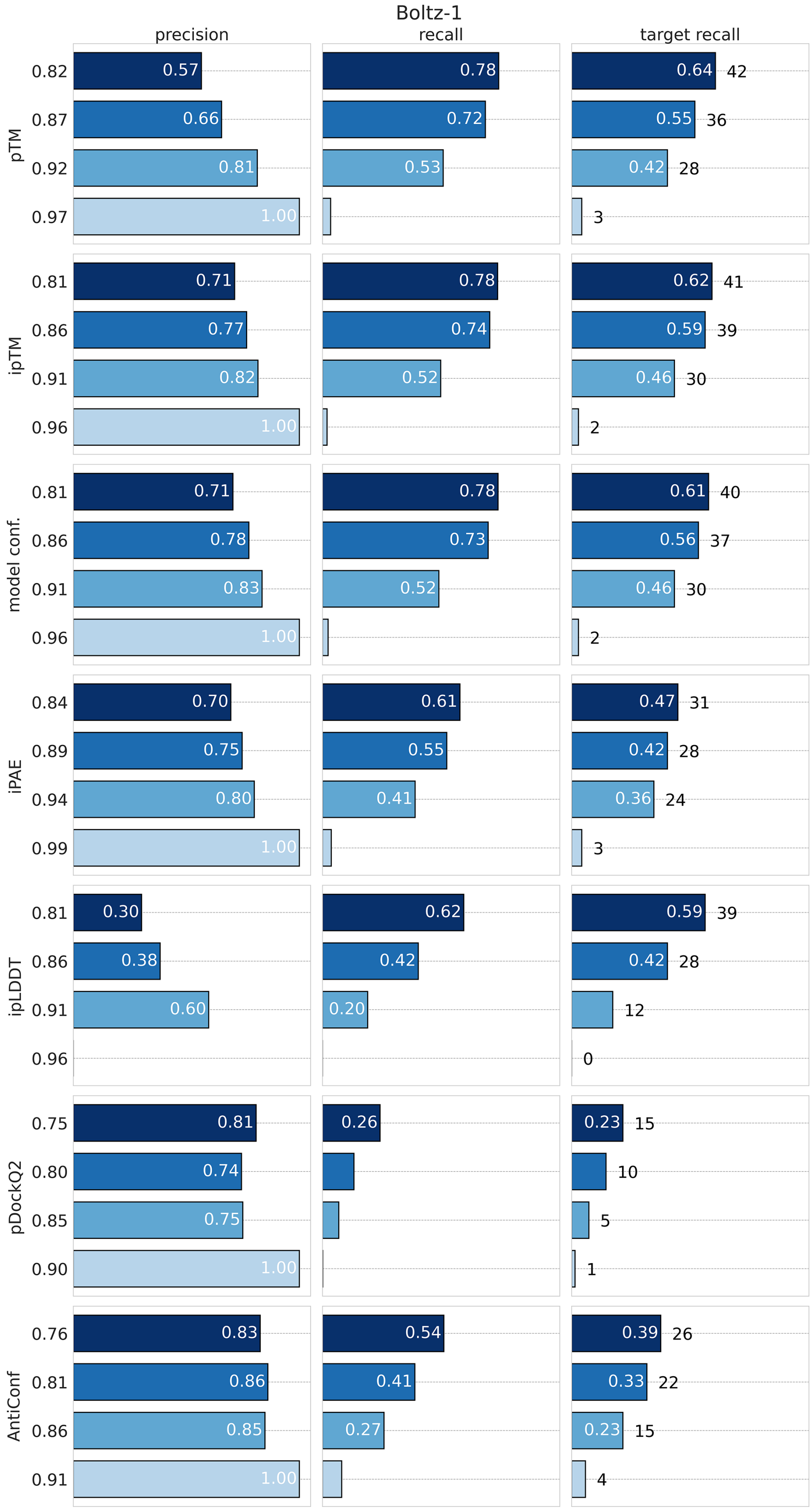

**Fig S9.** Boltz-1 performance using thresholds starting from the lowest value that achieves a precision of 1, and then decrementing it by 0.05, 0.10, and 0.15.

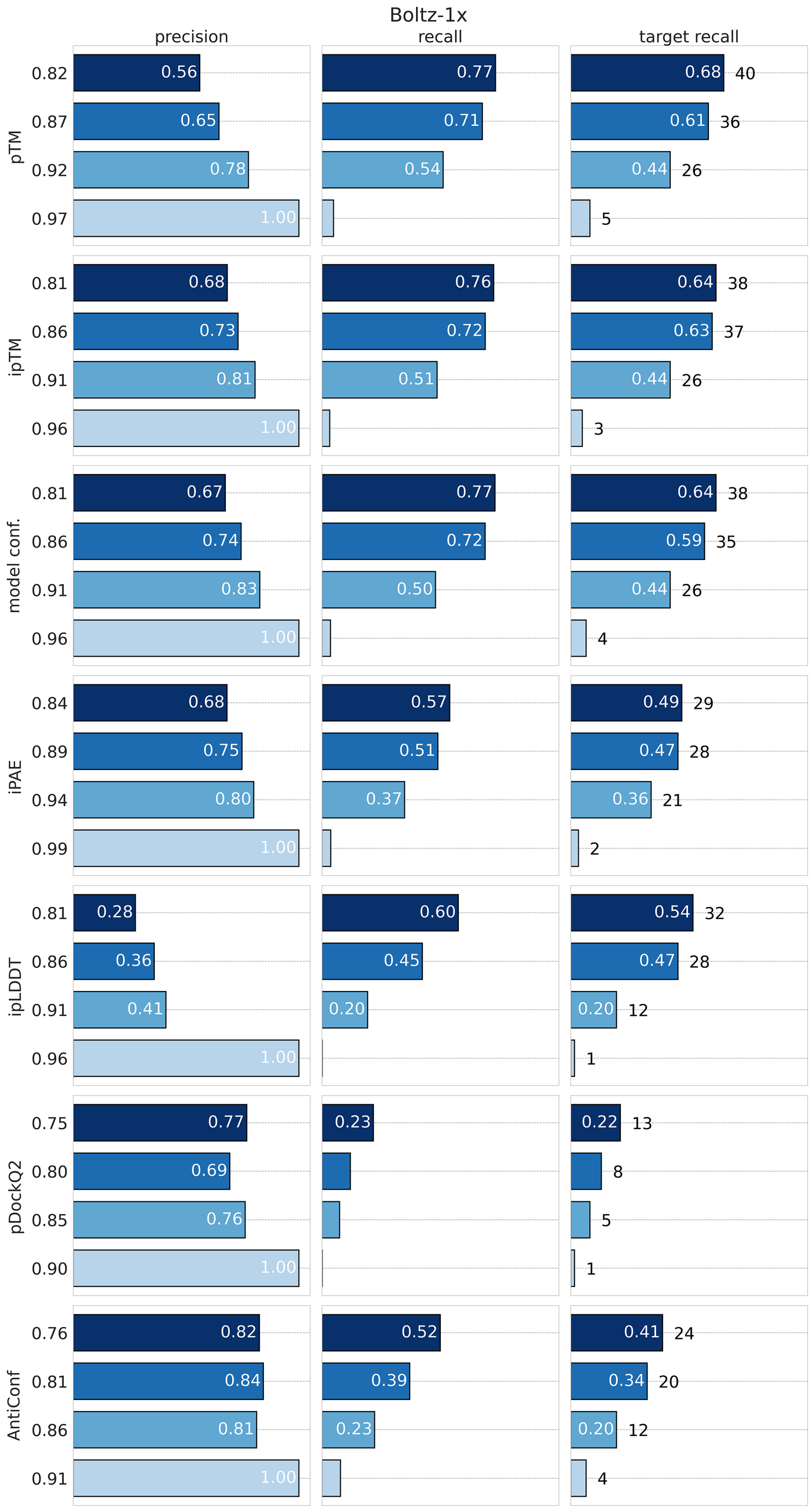

**Fig S10.** Boltz-1x performance using thresholds starting from the lowest value that achieves a precision of 1, and then decrementing it by 0.05, 0.10, and 0.15.

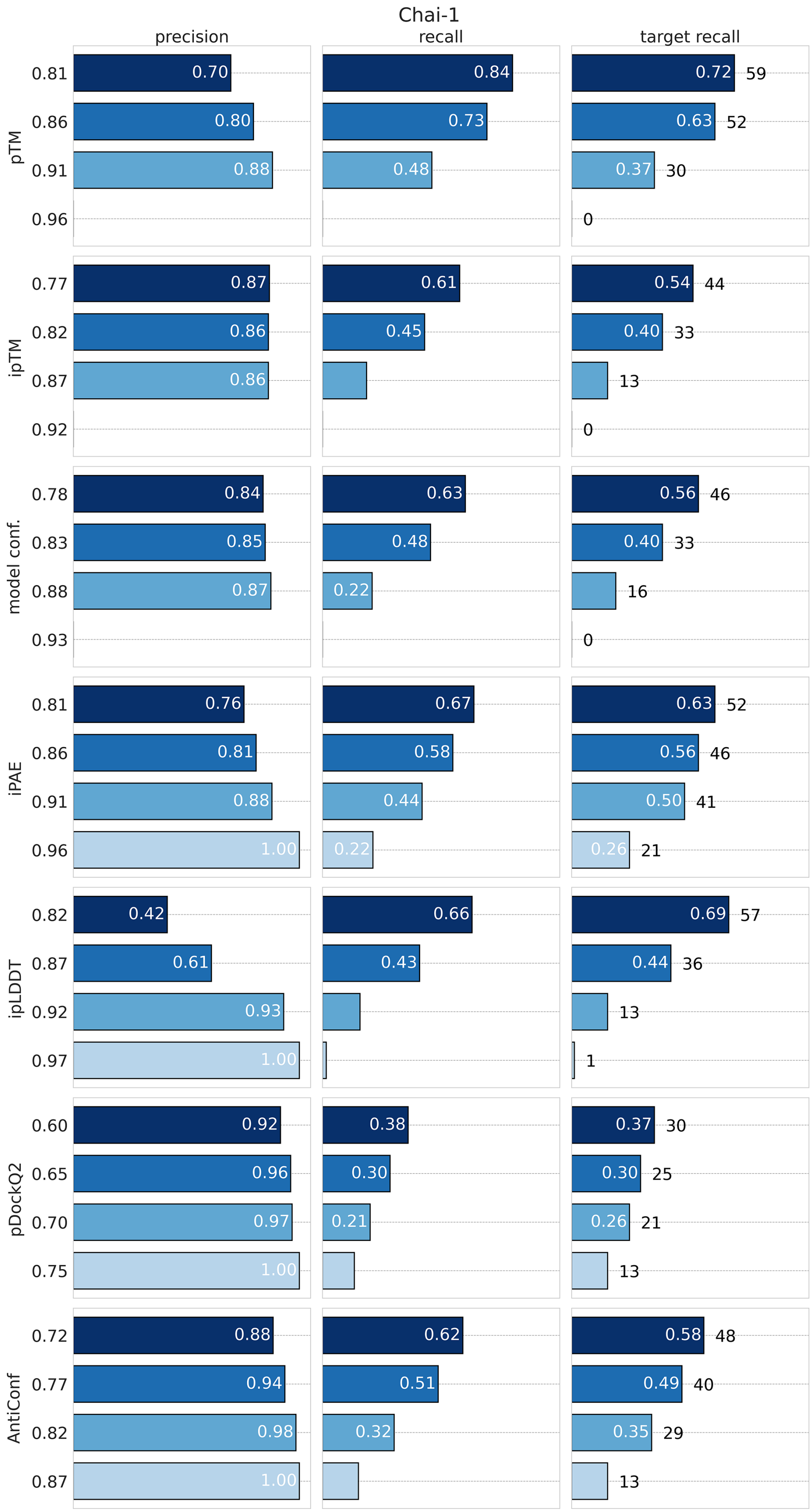

**Fig S11.** Chai-1 performance using thresholds starting from the lowest value that achieves a precision of 1, and then decrementing it by 0.05, 0.10, and 0.15

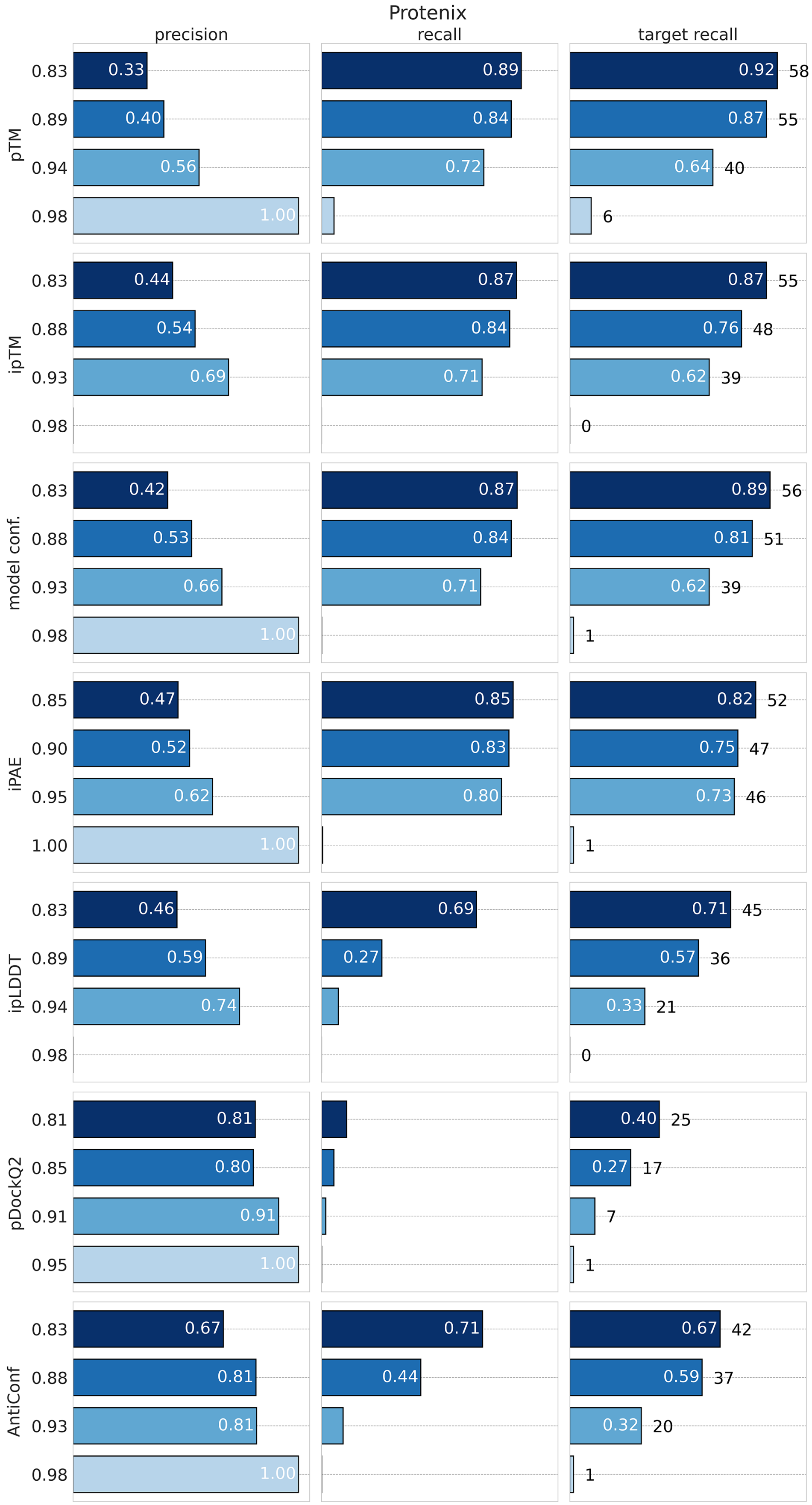

**Fig S12.** Protenix performance using thresholds starting from the lowest value that achieves a precision of 1, and then decrementing it by 0.05, 0.10, and 0.15.

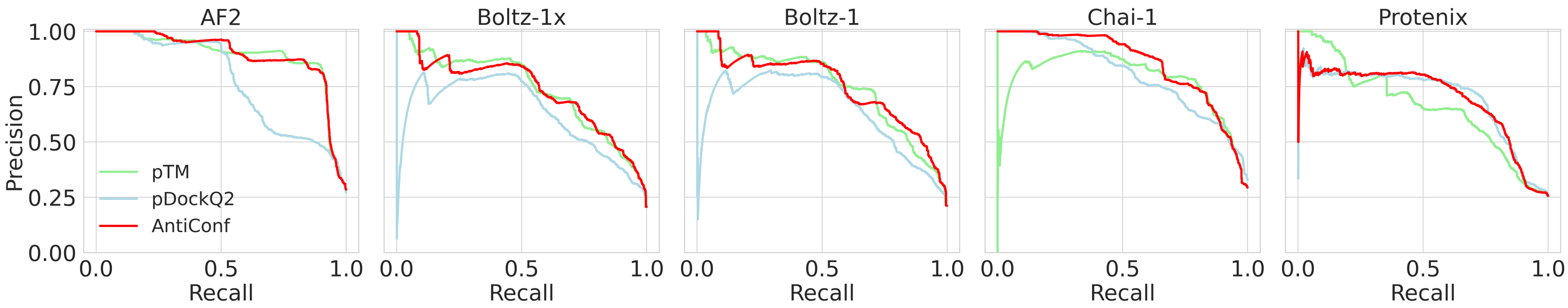

**Fig S13.** Precision-recall curves for pTM, pDockQ2 and AntiConf.
